# Intracellular lipopolysaccharide binds RETREG1/FAM134B to regulate ER remodeling upon bacterial infection

**DOI:** 10.1101/2024.07.25.605133

**Authors:** Yi-Lin Cheng, João Mello-Vieira, Adriana Covarrubias-Pinto, Alexis Gonzalez, Santosh Kumar Kuncha, Chun Kew, Kaiyi Zhang, Muhammad Awais Afzal, Nour Diab, Sophia Borchert, Siou-Ying Hong, Timothy Chun Huang, Wenbo Chen, Uxía Gestal Mato, Mathias Walter Hornef, Christian A. Hübner, Michael Hensel, Ivan Dikic

**Affiliations:** Institute of Biochemistry II, Faculty of Medicine, Goethe University, Frankfurt, Germany; Buchmann Institute for Molecular Life Sciences, Goethe University, Frankfurt, Germany; Institute of Basic Medical Sciences, College of Medicine, National Cheng Kung University, Tainan, Taiwan; Center of Infectious Disease and Signaling Research, National Cheng Kung University, Tainan, Taiwan; Altos Labs Inc, Bay Area Institute, Redwood City, CA, USA; Institute of Medical Microbiology, RWTH University Hospital, Aachen, Germany; Institute of Human Genetics, Jena University Hospital, Friedrich Schiller University, Jena, Germany; Abt. Mikrobiologie, Universität Osnabrück, Osnabrück, Germany; School of Medicine, College of Medicine, National Cheng Kung University, Tainan, Taiwan; Max Planck Institute of Biophysics, Frankfurt, Germany; Fraunhofer Institute for Molecular Biology and Applied Ecology (IME)

**Author notes:** Correspondence **Contact** Yi-Lin Cheng Institute of Basic Medical Sciences, College of Medicine, National Cheng Kung University, Tainan, Taiwan; Ivan Dikic Institute of Biochemistry II, Faculty of Medicine, Goethe University, Frankfurt, Germany.

**Keywords:** ER remodeling, lipopolysaccharide, outer membrane vesicles, RETREG1, *Salmonella*, xenophagy

## Abstract

Selective autophagy of the endoplasmic reticulum (ER), termed ERphagy or reticulophagy, plays a key role in organelle remodeling and cellular homeostasis. However, whether and how ERphagy is regulated during Gram-negative bacteria infection to influence host responses remains unclear. Here, we show that *Salmonella enterica* serovar Typhimurium releases lipopolysaccharide (LPS) that colocalizes with RETREG1/FAM134B, a reticulon-like ER-resident receptor for ERphagy. Cytosolic delivery of LPS, either during infection or via transfection, markedly increases RETREG1- and LC3B-decorated ER fragments. Mechanistically, affinity-isolation assays demonstrate that LPS directly binds RETREG1 through interactions between lipid A and positively charged residues within its amphipathic helices and C-terminal region. This interaction promotes RETREG1 oligomerization and drives ER membrane fragmentation, a process further amplified by the O-antigen moiety of LPS. The resulting ER fragments accumulate around LC3-positive *Salmonella*-containing vacuoles, facilitating bacterial clearance. Importantly, both intracellular and extracellular *Salmonella* exploit outer membrane vesicles (OMVs) to deliver LPS into the host cytosol, triggering RETREG1 activation and ER remodeling. Collectively, our findings reveal a previously unrecognized host response by which LPS of Gram-negative bacteria are sensed by the host ERphagy machinery to promote xenophagy and enhance antibacterial defense.

## Introduction

Multiple intracellular Gram-negative bacteria have been identified to manipulate host membrane system, including the endoplasmic reticulum (ER), which is the most abundant membranous system in mammalian cells [1]. However, whether there is a common component of Gram-negative bacteria participating in the regulation of ER membranes is unclear. The selective macroautophagy/autophagy of the ER (ERphagy) has recently been identified as the master mechanism for ER remodeling and turnover [2], enabling lysosomal degradation of different ER components, which are engulfed into autophagosomes through the interaction of MAP1LC3B/LC3B (microtubule associated protein 1 light chain 3 beta) with ERphagy receptors [2–4]. Eight mammalian ERphagy receptors have been identified, including the RETREG/FAM134 family [5–7], RTN3 [8], SEC62 [9], CCPG1 [10], ATL3 [11], TEX264 [12,13], and CALCOCO1 [14], which facilitate the degradation of different ER subdomains and respond to different types of stress conditions. The RETREG family and RTN3L, in particular, are able to induce significant ER remodeling. They contain two hairpin-like domains (transmembrane domains; TM) anchored into the lipid bilayer and connected by an amphipathic helix (AH) linker, as well as another carboxy-terminal AH, which is connected to the second TM. Together they form a reticulum homology domain (RHD), whose integrity appears critical for ERphagy [5,8,15]. The RHD of RETREG1, the best-studied ERphagy receptor, can sense highly positive membrane curvature, thereby triggering the formation of RETREG1 clusters that provide an additional force for ER membrane budding and scission [15–18].

LPS is the main component of the outer membrane of Gram-negative bacteria. It is composed of a conserved lipid A moiety, the inner core and outer core oligosaccharides, and O-antigen (O-Ag), the outermost part composed of a variable number of repeating sugar residues. In recent decades, most studies have focused on the extracellular role of LPS, which is specifically sensed on the plasma membrane by the TLR4 (toll like receptor 4)-LY96/myeloid differentiation factor 2 (lymphocyte antigen 96) co-receptor complex, that acts in conjugation with CD14 (CD14 molecule) to trigger signaling cascades for the expression of inflammatory genes [19]. The release of LPS by intracellular *Salmonella* to cytosolic vesicles was reported by Portillo and colleagues as early as 1997, however the targeted proteins and downstream effects are still elusive [20]. More recently, human CASP4 (caspase 4) and CASP5 have been identified as innate immune receptors that specifically recognize intracellular LPS [21]. The lipid A moiety of LPS directly binds to the amino (N)-terminal caspase-activation and recruitment domain of these caspases, triggering their oligomerization and activation [21]. Activated CASP4 further cleave GSDMD (gasdermin D) to liberate its N-terminal domain, which has membrane-pore-forming activity at the plasma membrane and thereby induces pyroptosis [22,23]. Moreover, LPS on cytosolic *Salmonella* is also recognized by GBP1 (guanylate binding protein 1), which initiates the assembly of a GBP coat for CASP4 recruitment and activation to trigger non-canonical inflammasome signaling [24].

Extracellular Gram-negative bacteria also expose LPS to the cytosol, by having their LPS-enriched outer membrane vesicles (OMVs) internalized into the cell [25]. The bacterial envelop is composed of the cytoplasmic membrane and the outer membrane, which enclose the periplasmic space containing a peptidoglycan layer and periplasmic proteins that stabilize the membrane through various crosslinks [26–28]. Upon infection, environmental stress factors released by the host disturb the stability of those crosslinks and increase outer membrane budding into OMVs with a diameter of 20 nm to 250 nm. OMVs are internalized into the host cell via endocytosis, and LPS is released from early endosomes into the cytosol, where it activates CASP4/caspase-11 [25] and possibly exerts other effects.

Here we describe a novel function for intracellular LPS mediated by binding to the RETREG1 ERphagy receptor. Both *Salmonella* infection and intracellular LPS induced RETREG1-mediated ER fragmentation. *Salmonella* LPS colocalized with RETREG1 as determined by immunofluorescence experiments, and biochemical analysis showed that lipid A of LPS directly bound to RETREG1 via electrostatic forces. Although lipid A was sufficient for RETREG1 binding, oligomerization of RETREG1 and the following membrane fragmentation required the presence of O-Ag of LPS. RETREG1-associated ER membrane then surrounded xenophagosomes promoting bacterial clearance. We also found that OMVs are responsible for the delivery of LPS from both intracellular and extracellular bacteria into the cytosol. Taken together, this study identified RETREG1 as an intracellular sensor of LPS, enabling the ER membrane remodeling and xenophagy.

## Results

### LPS colocalizes with RETREG1 and triggers RETREG1-mediated ER fragmentation

To investigate whether intracellular Gram-negative bacteria activate ERphagy, we first examined the role of the ERphagy receptor RETREG1 during *Salmonella enterica* serovar Typhimurium SL1344 infection. In HeLa TREx FLAG-HA-RETREG1-inducible cells, *Salmonella* infection strongly induced colocalization of RETREG1 puncta with LC3B puncta, indicating induction of RETREG1-mediated ER fragmentation upon infection (Figure 1A). Comparing to RETREG1, RTN3 did not show significant colocalization with LC3B after *Salmonella* infection (Figure S1A). To our surprise, we found LPS puncta colocalizing with RETREG1 which were not associated with bacteria, indicated by nucleic acid staining with 4’,6-diamidino-2-phenylindole (DAPI), either in wild-type (WT) RETREG1 or LC3B-interacting region mutant (mtLIR) RETREG1 (Figure 1A and Figure S1B). Moreover, endogenous RETREG1, also colocalized with *Salmonella* LPS after infection (Figure S1C). High throughput analysis by confocal imaging cytometer CQ1 showed that around 20% of LPS puncta colocalized with RETREG1 after *Salmonella* infection (Figure 1B). Despite most of RETREG1 puncta colocalize with LPS or LC3B, respectively (arrows), some of the RETREG1 puncta colocalize with LPS and LC3B simultaneously (arrowheads) as the histogram showed (Figure 1A). The number of RETREG1- and LC3B-double-positive puncta after *Salmonella* infection was significantly increased in WT RETREG1-expressing HeLa cells, but not in mtLIR RETREG1-expressing HeLa cells, indicating *Salmonella* infection caused induction of RETREG1-mediated ERphagy and ER fragmentation (Figure 1C and Figure S1D). Co-immunoprecipitation (co-IP) assays revealed that *Salmonella* infection enhanced interaction of RETREG1 with LC3B-II in the presence or absence of bafilomycin A_1_ (BafA1), which inhibits maturation of autophagosomes for lysosomal degradation (Figure 1D). Our previous study defined ER fragments as puncta containing RETREG1, LC3B-II and ER proteins that do not directly bind LC3B [18]. To clarify whether LPS can trigger RETREG1-mediated ER fragmentation, we transfected *Salmonella* LPS in HeLa cells followed by immunostaining and confocal microscopy analysis. The results showed that LPS colocalizes with RETREG1 puncta and LC3B puncta simultaneously (arrowheads) showing in the images and the histogram (Figure 1E). The number of RETREG1- and LC3B-double-positive puncta also increased in presence of LPS (Figure 1F). Moreover, transfection of *Salmonella* LPS triggered RETREG1 fragments delivered to lysosomes, indicated by LAMP1 (Figure S1E). Thus, *Salmonella* LPS initiated RETREG1-mediated ER fragmentation.

**Figure 1.**
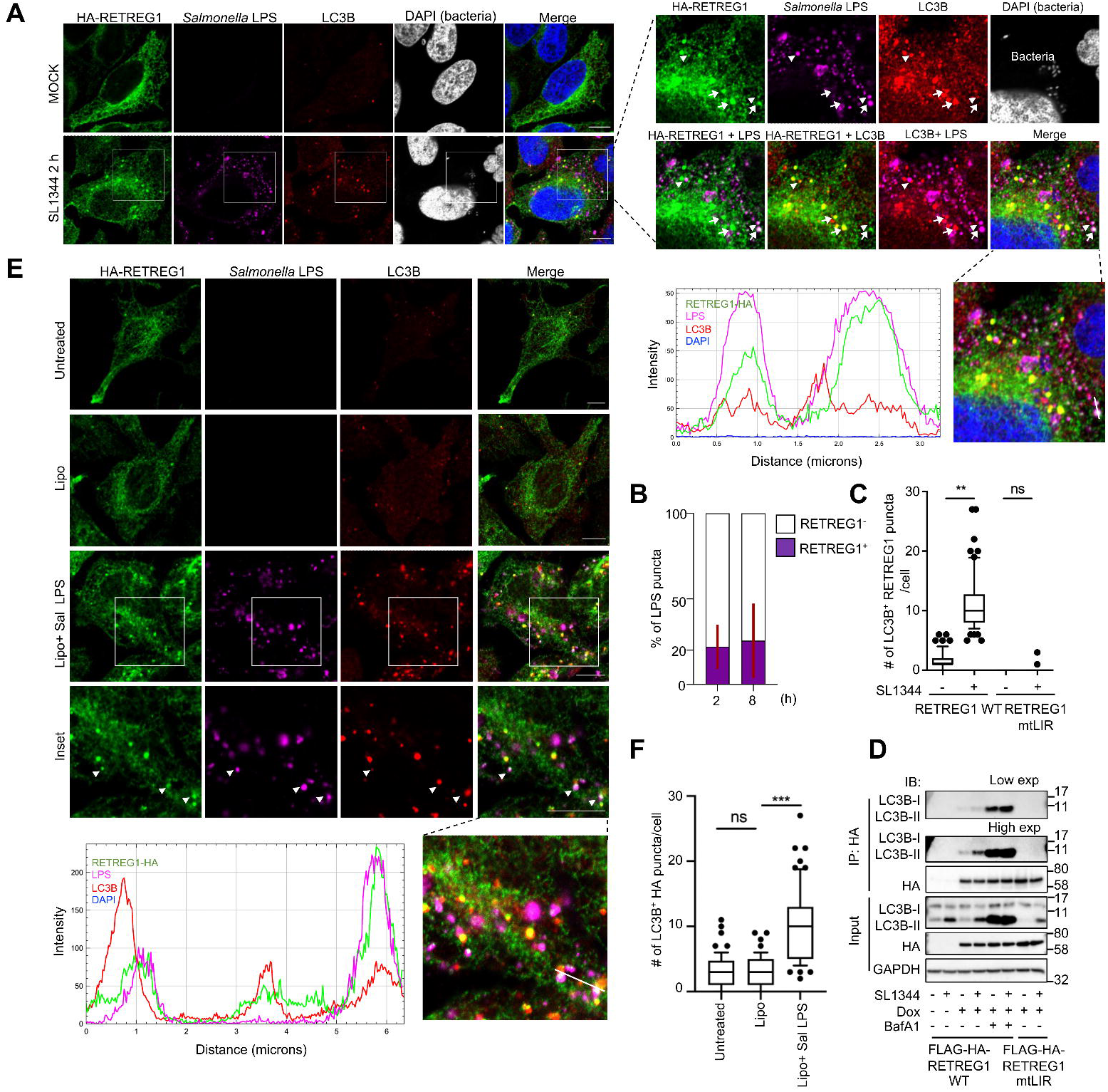
*Salmonella* LPS activates RETREG1-mediated ER fragmentation. (**A**) Immunofluorescence staining of HA, *Salmonella* LPS, and LC3B in HeLa TREx inducible cell lines expressing FLAG-HA-RETREG1 that were infected with SL1344 for 2 h. DAPI staining was used for cell nuclear and bacteria DNA staining. Arrows indicate the puncta with colocalization of HA-RETREG1 and *Salmonella* LPS or LC3B, and arrowheads indicate the puncta with colocalization of HA-RETREG1, *Salmonella* LPS and LC3B. Scale bar: 10 μm. The histogram showed the florescence intensity and the location of HA-RETREG1 (green), LC3B (red), and *Salmonella* LPS (magenta) of the white line in the image, which analyzed by image J. (**B**) Quantification of HA and *Salmonella* LPS staining in HeLa TREx inducible cell lines expressing FLAG-HA-RETREG1 infected with SL1344 for 2 h and 8 h. The percentage of RETREG1^+^ and RETREG1^+^ LPS puncta in all LPS puncta were quantified using CellPathfinder software. Data presented as mean + standard error of the mean (sem), n=3, > 400 cells. (**C**) Number of LC3B^+^ HA-RETREG1 puncta in HeLa TREx inducible cell lines expressing wild type (WT) or LIR mutant (mtLIR) FLAG-HA-RETREG1 infected with SL1344 for 2 h. Representative images are shown in Figure S1D. (**D**) Co-IP of endogenous LC3B-II with HA in HeLa TREx inducible cells without doxycycline (Dox) treatment or with FLAG-HA-RETREG1 WT or FLAG-HA-RETREG1 mtLIR after Dox treatment. The cells were infected with SL1344 for 2 h and pretreated with 200 ng/ml bafilomycin A_1_ (BafA1) for 3 h. GAPDH was used as an internal control of input. (**E** and **F**) Immunofluorescence staining and quantification of HA, *Salmonella* LPS, and LC3B in HeLa TREx inducible cell lines expressing FLAG-HA-RETREG1 transfected with 1 μg/ml *Salmonella* LPS for 18 h using lipofectamine 2000. Lipo, lipofectamine only. (**E**) Arrowheads indicate the puncta with colocalization of HA-RETREG1, *Salmonella* LPS and LC3B. The histogram showed the florescence intensity and the location of HA-RETREG1 (green), LC3B (red), and *Salmonella* LPS (magenta) of the white line in the image, which analyzed by imageJ. Scale bar: 10 μm. (**F**) Quantification of LC3B^+^ HA-RETREG1 puncta per cell. (**C** and **F**) Solid bars of boxes indicate the medians. Boxes represent interquartile range from 25^th^ to 75^th^ percentile, and whiskers indicates 10^th^ to 90^th^ percentile. Differences were statistically analyzed by one-way ANOVA and Tukey’s multiple comparison test. Data were collected 60 cells from three independent biological replicates. **p < 0.01, ***p < 0.001. ns, not significant.

### LPS binds to RETREG1 via electrostatic force

Since LPS colocalized with RETREG1 in the confocal images, we hypothesized that LPS binds directly to RETREG1. Therefore, we carried out a series of streptavidin pull down assays to clarify the mode of interaction (Figure 2A). We used biotin-conjugated *E. coli* O111:B4 LPS immobilized on streptavidin sepharose to study its interaction with either HA-RETREG1 or HA-RTN3L (Figure 2B). We detected that biotin-LPS strongly pulled down HA-RETREG1 in a dose-dependent manner, while contained much less affinity to HA-RTN3L (Figure 2B). The endogenous RETREG1 in the lysates of HeLa cell also was pulled downed by biotin-LPS strongly compared to the other ERphagy receptors, RTN3L and TEX264 (Figure 2C). The binding of LPS to RETREG1 was LIR motif-independent (Figure S2A). In addition to *in vitro* pulldown assays, pulling down the biotin-LPS which delivered into cells expressing FLAG-HA-RETREG1 or FLAG-HA-RTN3L also showed strong interaction to RETREG1 (Figure S2B). Upon *Salmonella* infection in cells expressing FLAG-HA-RETREG1, immunoprecipitation of HA-RETREG1 was also able to detect the interaction of LPS, but no significant interaction between RTN3L and LPS was detected upon *Salmonella* infection (Figure 2D). To further clarify which segment of LPS is required for binding to RETREG1, we performed a competition assay. WT LPS (also called smooth [S]-LPS), rough LPS, including Ra-LPS (LPS without O-Ag) and Re-LPS (LPS containing lipid A and three 2-keto-3-deoxy-octonate [KDO]), and lipid A (Figure 2E), were preincubated with lysates containing FLAG-HA-RETREG1, followed by incubation with biotin-LPS-immobilized streptavidin sepharose. We analyzed whether the preincubation with different LPS variants can block the binding of biotin-LPS to RETREG1 (Figure 2F). The results revealed that all kinds of LPS significantly reduced the binding ability of biotin-LPS to RETREG1, except muramyl dipeptide (MDP), which is another component of the bacteria cell wall, demonstrating that lipid A is sufficient for RETREG1 binding (Figure 2F). Furthermore, polymyxin B, a cationic peptide antibiotic, which interacts with lipid A via ionic and hydrophobic forces, inhibited the binding of biotin-LPS to RETREG1 in a dose-dependent manner (Figure 2G).

**Figure 2.**
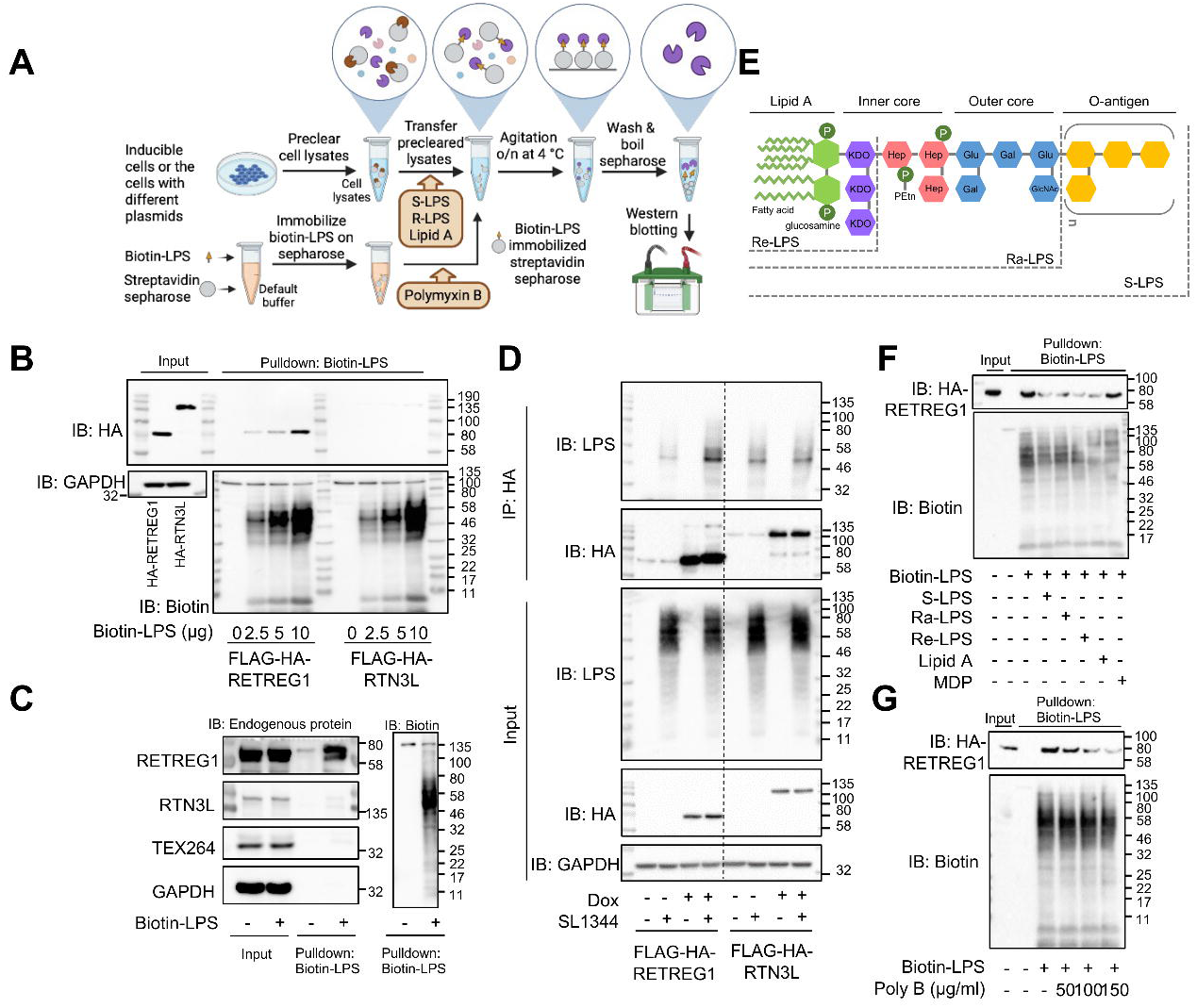
LPS binds to RETREG1 via lipid A moiety. (**A**) Schematic representation of streptavidin affinity-isolation assays. (**B**) Streptavidin pulldown assays that determine the binding of the indicated amounts of biotin-conjugated *E. coli* O111:B4 LPS and HA-tagged RETREG1 or RTN3L in FLAG-HA-RETREG1- or FLAG-HA-RTN3L-expressing HeLa TREx cell lysates. GAPDH was used as an internal control of input. (**C**) Streptavidin pulldown assays that determine the binding of the biotin-conjugated *E. coli* O111:B4 LPS to the endogenous protein of the ERphagy receptors, RETREG1, RTN3L and TEX264 in HeLa cell lysates. GAPDH was used as an internal control of input. (**D**) Co-IP of LPS with HA in HeLa TREx inducible cell lines without Dox treatment or expressing FLAG-HA-RETREG1 or FLAG-HA-RTN3L by Dox treatment, which were infected with SL1344 for 2 h. GAPDH was used as an internal control of input. (**E**) Schematic representation of the LPS structure and the segments of different mutant variants of LPS, including lipid A, Re-LPS, Ra-LPS, and smooth (S)-LPS. P, phosphate; KDO, 3-deoxy-D-*manno*-octulosonic acid; Hep, L-*glycero*-D-*manno*-heptose; PEtn, phosphatidyl-ethanolamine; Glc, D-glucose; Gal, D-galactose; GlcNac, N-acetyl-D-glucosamine. (**F**) Unlabeled S-LPS, Ra-LPS, Re-LPS, lipid A, and muramyl dipeptide (MDP) were used for competition with the binding of biotin-conjugated *E. coli* O111:B4 LPS to HA-tagged RETREG1 in Dox-induced FLAG-HA-RETREG1 HeLa TREx cell lysates. (**G**) Indicated concentrations of polymyxin B (Poly B) were used to inhibit the binding of biotin-conjugated *E. coli* O111:B4 LPS to HA-tagged RETREG1 in Dox-induced FLAG-HA-RETREG1 HeLa TREx cell lysates.

Next, we characterized how RETREG1 binds to LPS. Since lipid A contains multiple negatively charged groups [29], we hypothesized that the interaction might occur through electrostatic forces. We analyzed each domain of RETREG1 and found that two cytoplasmic amphipathic helices, the linker and C-terminal region of the amphipathic helices (AHL and AHC), and the C-terminal region (Cterm), are highly solvent-accessible and enriched with positively charged residues, including arginine (R) and lysine (K) (Figure 3A). According to this analysis, we mutated R or K residues to glutamic acid (E) in these three RETREG1 regions (Figure 3B). We then evaluated the interactions of these mutants with biotin-LPS.

**Figure 3.**
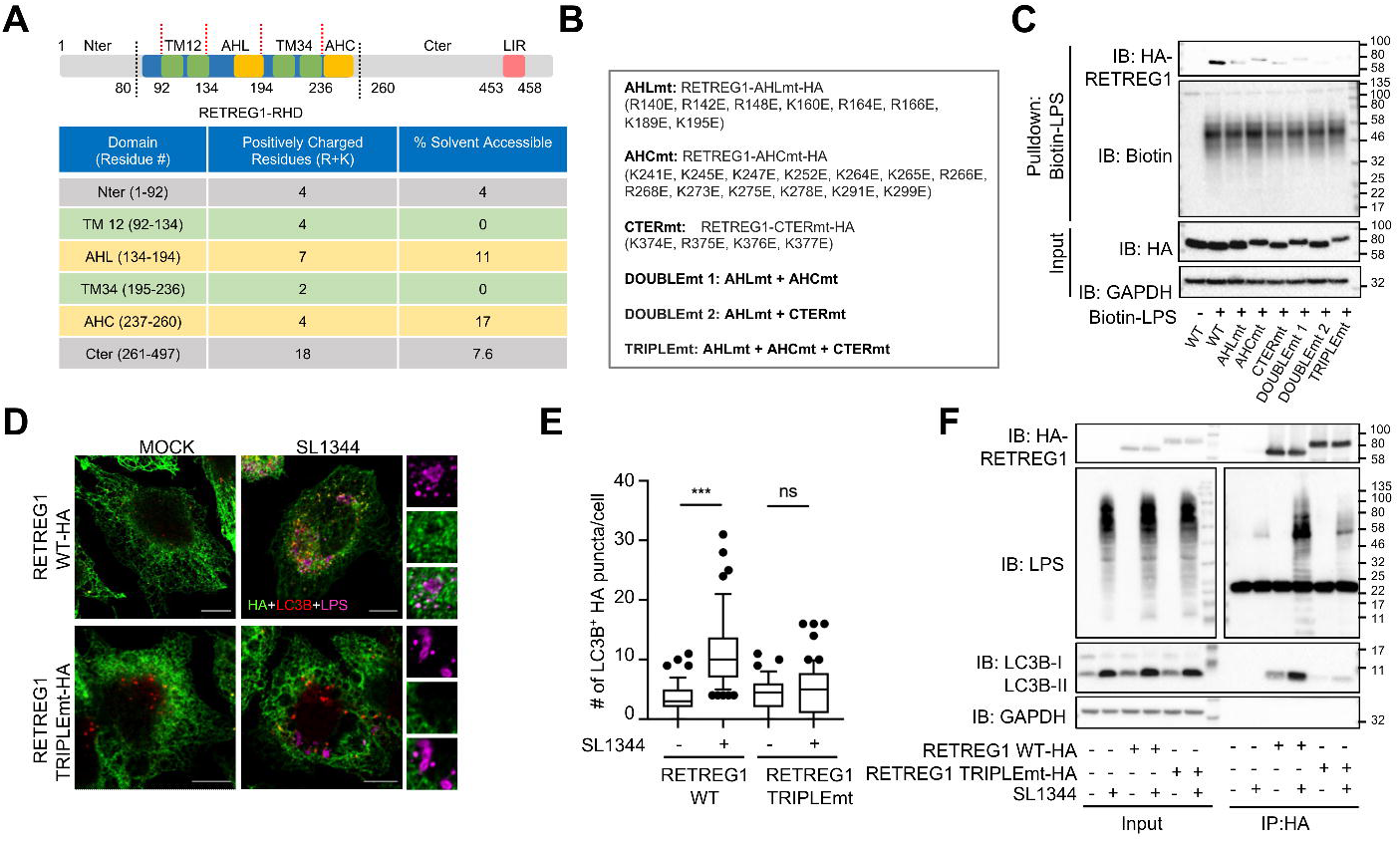
RETREG1 binds to LPS via electrostatic forces. (**A**) Schematic representation of the full length RETREG1 sequence, which is composed of a reticulum homology domain (RHD), N-terminal (Nter, gray) and C-terminal regions (Cter, grsy). LC3-interacting region consisting of 4 amino acids (LIR, pink) is located in the C-terminal region. The RHD consists of two transmembrane segments TM12 and TM34 (green), which are connected by a 60-residue linker, and two additional terminal segments. The C-terminal fragment of the RHD (AHC, yellow) and the linker-helix (AHL, yellow) form conserved amphipathic helices. The lower table shows the number of positively charged residues (arginine, R; Lysine, K) and the percentage of solvent accessible in every segment of RETREG1. (**B**) Substitution of amino acids (R or K to glutamic acid, E) in segment AHL (AHLmt), AHC (AHCmt), and C-terminal (CTERmt). DOUBLEmt 1, DOUBLEmt 2, and TRIPLEmt indicate combination as shown in the table. (**C**) Streptavidin pulldown assays that determine the binding of the indicated amount of biotin-conjugated *E. coli* O111:B4 LPS to HA-tagged wild type or different mutant RETREG1 transfected in HEK293T cells. GAPDH was used as an internal control of input. (**D** and **E**) Immunofluorescence staining and quantification of HA and *Salmonella* LPS, and LC3B in RETREG1 WT-HA or RETREG1 TRIPLEmt-HA transfected HeLa cells infected with SL1344 for 2 h. Scale bar: 10 μm. (**E**) Quantification of number of LC3B^+^ HA-RETREG1 puncta per cell. Solid bars of boxes indicate the medians. Boxes represent interquartile range from 25^th^ to 75^th^ percentile, and whiskers indicates 10^th^ to 90^th^ percentile. Differences were statistically analyzed by one-way ANOVA and Tukey’s multiple comparison test. ***p < 0.001. ns, not significant. Data were collected 60 cells from three independent biological replicates. (**F**) Co-IP of LPS and endogenous LC3B-II in RETREG1 WT-HA or RETREG1 TRIPLEmt-HA transfected HeLa cells, which were infected with SL1344 for 2 h. GAPDH was used as an internal control of input. The bands at ∼ 22 kDa is the light chain of the antibody used for co-IP.

Each mutant RETREG1 construct showed significantly reduced binding to biotin-LPS, and binding was completely abolished when mutations in all three regions were combined (Figure 3C). Transfection of RETREG1 TRIPLEmt (mutations in all three regions) robustly blocked colocalization of RETREG1 with LPS and LC3B (Figure 3D, E). These results were also confirmed by co-IP experiments in which RETREG1 TRIPLEmt-HA co-precipitated significantly less LPS and LC3B-II compared to RETREG1 WT-HA (Figure 3F). Moreover, *in vitro* streptavidin affinity-isolation assays showed that biotin-LPS pulled down RETREG3/FAM134C and RETREG1 with similar efficiency whereas RETREG2/FAM134A interacted significantly weaker (Figure S2C). This result could be explained by the fact that RETREG1 and RETREG3 have a high sequence similarity, from which the sequence of RETREG2 clearly deviates (Figure S2D, E, F).

### RETREG1 oligomerization is triggered by O-Ag of LPS

Oligomerization of RETREG1 is reported to drive ER membrane scission followed by ERphagy induction [17]. Especially, caspases also undergo oligomerization after binding with LPS [21]. We therefore hypothesized that LPS binds to RETREG1 to induce its oligomerization. S-protein-FLAG-streptavidin binding peptide (SFB)-RETREG1 was expressed and purified from HEK293T cells followed by incubation with S-LPS or lipid A. In order to analyze oligomeric species of RETREG1, we performed blue native PAGE, which enforces migration of native protein complexes or oligomers based solely on their molecular weight [30], followed by immunoblotting with FLAG. S-LPS significantly increased the presence of RETREG1 oligomers of approximately 720 kDa (Figure 4A). To our surprise, lipid A did not show a similar effect, although it played a major role in binding to RETREG1, suggesting that O-Ag is indispensable for the oligomerization of RETREG1. To test this idea, we infected HeLa cells expressing SFB-RETREG1 with an isogenic set of *Salmonella* mutant strains with defined defects in the O-antigen biosynthesis, including the *rfbP* (O-antigen transferase) and *rfaL* (O-antigen ligase) mutant strains lacking O-Ag, the *rfaG* (glucosyltransferase I) mutant strain lacking O-Ag and outer core, as well as the *rfaH* (transcriptional activator) mutant strain with reduced polymeric O-Ag (Figure 4B, C) [31]. After infection, SFB-RETREG1 was purified, and blue native PAGE analysis was performed. The results showed that RETREG1 oligomers in the range of ∼720 kDa to ∼1048 kDa were significantly reduced after infection with any of the O-Ag-defective strains compared to WT and complemented strains (Figure 4B).

**Figure 4.**
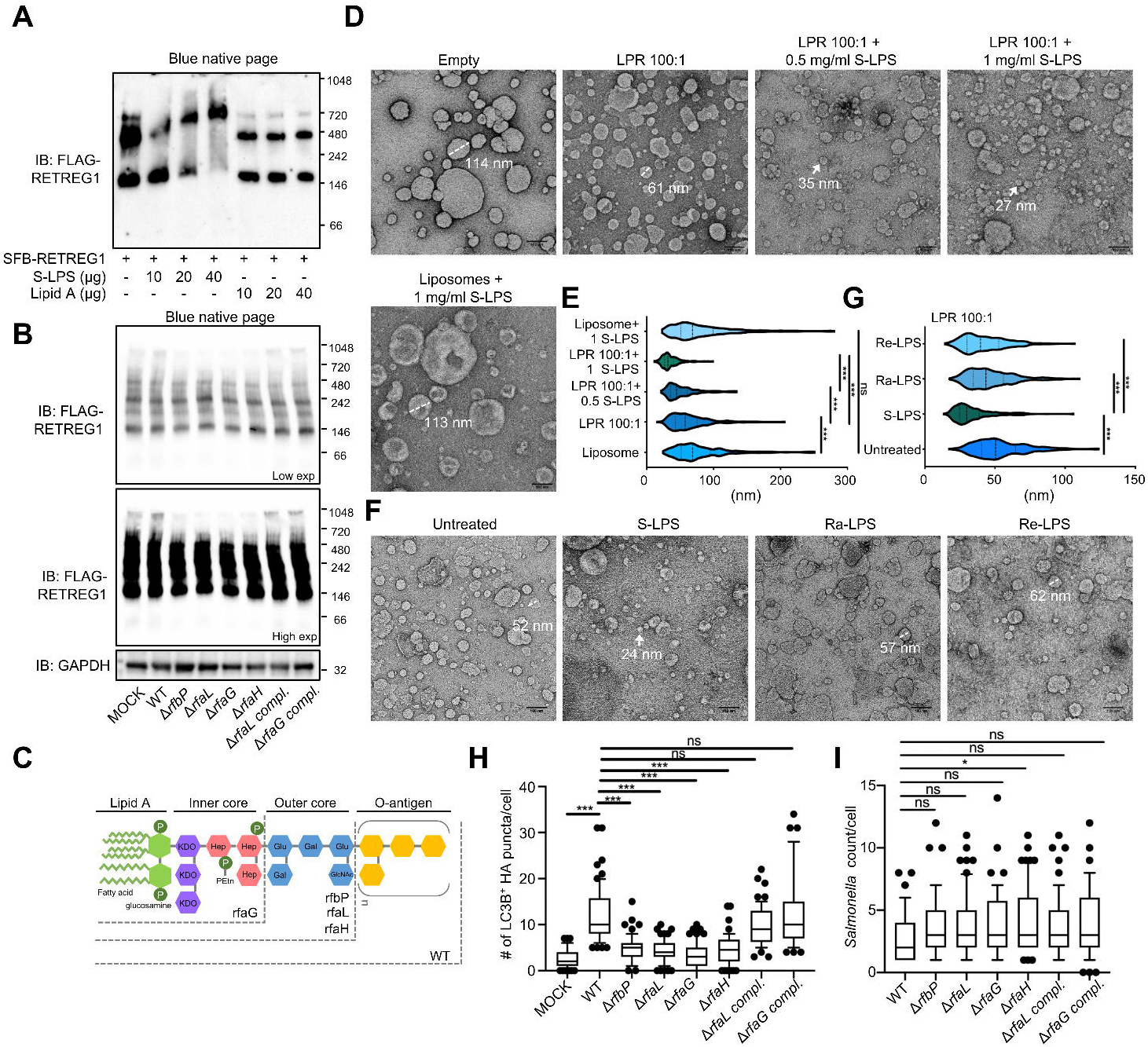
O-Ag of LPS induces RETREG1 oligomerization and membrane fragmentation. (**A**) Size analysis of S protein-FLAG-Streptavidin binding peptide (SFB)-RETREG1 oligomers after incubation with indicated amount of *Salmonella* S*-*LPS or lipid A by blue native page and immunoblotting of FLAG. (**B**) Size analysis of SFB-RETREG1 oligomers purified from cell lysates infected with NCTC12023 WT or O-Ag mutant (Δ) and complemented (compl.) strains. Upper panel shows lower exposure (Low exp) blot and lower blot shows higher exposure (high exp) blot. GAPDH was used as an internal control of input. (**C**) Schematic representation of LPS structure of different O-Ag mutant strains, including the strains mutant for *rfbP, rfaL*, *rfaG,* and *rfaH.* (**D** and **E**) Representative images (**D**) and quantitative results (**E**) of remodeled proteoliposomes (scale bars, 100 nm) taken from negative stain Transmission Electron Microscopy (nsTEM). Empty liposomes (Empty) were added RETREG1-GST recombinant protein at a lipid to protein ratio (LPR) 100:1 without (LPR 100:1) or with 0.5 mg/ml LPS (LPR 100:1+LPS 0.5) or 1 mg/ml LPS (LPR 100:1+ LPS 1). Incubation of liposome with 1 mg/ml LPS was taken as negative control to check the effect of LPS on liposome. The diameter of the representative proteoliposome with average size for each group showed by dotted line or arrow. (**F** and **G**) Representative images (**F**) and quantitative results (**G**) of different LPS, including 1 mg/ml of S-LPS, Ra-LPS, and Re-LPS, remodeled proteoliposomes (LPR 100:1, scale bars, 100 nm). The diameter of the representative proteoliposome with average size for each group showed by dotted line or arrow. (**E** and **G**) The size distributions (n=300 each) from nsTEM images were measured and showed by violin plots, showing a central boxplot (black lines indicate median with interquartile range) with mirrored histograms on both sides (**G**). (**H** and **I**) Immunofluorescence of HA, *Salmonella* LPS, and LC3B in HeLa TREx inducible cell line expressing FLAG-HA-RETREG1 infected with NCTC12023 WT or LPS O-Ag mutant (Δ) and complemented (compl.) strains for 2 h (**H**). DAPI staining was used for cell nuclear and bacterial DNA staining. Scale bar: 10 μm. Quantification of LC3B^+^ HA-RETREG1 puncta per cell (**H**) and *Salmonella* count per cell (**I**). Data were collected 60 cells from three independent biological replicates. Representative images are shown in **Figure S3C**. (**E**,**G**,**H**,**I**) Differences were statistically analyzed by one-way ANOVA and Tukey’s multiple comparison test. *p<0.05, **p<0.01, ***p < 0.001, ns not significant.

RETREG1 clustering would further induce membrane curvature and remodeling [15]. To further prove that O-Ag regulates membrane remodeling driven by RETREG1, we conducted a liposome-remodeling assay in which incubation with purified RETREG1 drives membrane fragmentation [15]. We first determined the suitable lipid-to-protein ratio (LPR) for the examination of the effect of LPS on RETREG1-triggered membrane fragmentation (Figure S3A, B). LPR 40:1 triggered strong liposome remodeling, indicated by a significant reduction of the liposome diameter compared to the control group of liposomes only (Empty). LPR 100:1 had a moderate effect, resulting in a slightly reduced liposome size (Figure S3A, B). Thus, we used LPR 100:1 to study the action of LPS (Figure 4D, E, F, G). S-LPS significantly enhanced the membrane remodeling effect of RETREG1 compared to liposomes incubated with RETREG1 alone in a dose-dependent manner (Figure 4D, E). Notably, LPS alone did not change the size of liposomes compared to the control sample, demonstrating that LPS cannot fragment membrane by itself, but enhances the action of RETREG1 on membranes (Figure 4D, E). Compared to S-LPS, Ra-LPS and Re-LPS, which lack the O-Ag, had less pronounced effects on the size of liposomes, i.e. on the ability of RETREG1 to induce membrane fragmentation (Figure 4F, G). To further verify the role of O-Ag in ER fragmentation in cells, we performed co-localization analysis of HA-RETREG1 and LC3B after infection of WT, the isogenic mutant strains without O-Ag, and the two complemented strains of Δ*rfaL* and Δ*rfaG* (Figure 4H, S3C). The number of RETREG1- and LC3B-double-positive puncta was significantly reduced upon infection with O-Ag mutant strains, Δ*rfbP,* Δ*rfaL,* Δ*rfaG,* and Δ*rfaH* strains, in comparison to the WT strain. On contrary, complemented strains of Δ*rfaL* and Δ*rfaG* induced a similar number of RETREG1- and LC3B-double-positive puncta as the WT strain (Figure 4H, S3C). The bacteria count per cell did not show significant differences among the groups at this early time point, implying the reduction in RETREG1- and LC3B-double-positive puncta was not due to fewer bacteria (Figure 4I, S3C). Compared to S-LPS, transfection of Ra-LPS, Re-LPS, and lipid A, which are the LPS without O-Ag, did not increase the number of HA-RETREG1- and LC3B-double-positive puncta (Figure S3D, E). The other two cell wall components of bacteria, Pam3CSK4 and MDP, did not affect puncta formation (Figure S3D, E). These results demonstrated that O-Ag of LPS triggered oligomerization of RETREG1 and RETREG1-mediated membrane remodeling.

### RETREG1 is recruited to xenophagosomes and promotes bacterial clearance

To further investigate the fate and function of RETREG1 after infection, we measured the RETREG1-mediated ERphagy flux by mCherry-EGFP-RETREG1 WT or LIR mutant-inducible HeLa cells after *Salmonella* infection for 2 h and 4 h (Figure S4A, S4B). Owing to the instability of EGFP in acidified environment, only the red fluorescence of mCherry would be detected when RETREG1 is delivered to lysosomes [5,13,17]. *Salmonella* infection strongly induced formation of red RETREG1 puncta 4 h post-infection in mCherry-EGFP-RETREG1 WT HeLa cells, but not in cells expressing the LIR mutant (Figure S4A, S4B). Moreover, RETREG1 which is positive for both EGFP and mCherry forming vacuoles surround bacteria (Figure S4A-1). In *Salmonella*-infected intestinal tissue sections of 5 days-old suckling mice, RETREG1 was also recruited to *Salmonella* (Figure S4C). The RETREG1-positive *Salmonella*-containing compartments were positive for ER markers, CANX and REEP5 as well (Figure S4D).

According to previous studies, invasive *Salmonella* can survive in the endolysosomal membrane-derived *Salmonella*-containing vacuoles (SCVs), but some populations of bacteria escape into cytosol, which can be cleared after sequestration within xenophagosomes or become hyperproliferating bacteria [32–36]. We decided to study whether the RETREG1-positive structures contribute to one of these two compartments. We observed that bacteria decorated with RETREG1-HA, can colocalize with LC3B but not with the endolysosomal marker LAMP1 (Figure 5A). The imaging analysis showed the percentage of LC3B-positive and RETREG1-positive *Salmonella* are similar, and both significantly accumulated after Baf A1 treatment, indicating these vacuoles underwent lysosomal degradation (Figure 5B). LAMP1-positive SCVs accounts around 40% of total *Salmonella*, which did not change obviously after Baf A1 treatment at 2 h post-infection, and even reduced at 4 h post-infection (Figure 5B). The imaging analysis further showed almost 60% LC3B-positive *Salmonella*-containing xenophagosomes were RETREG1-positive after *Salmonella* infection for 2 h, and around 40% at 4 h post-infection; conversely very few SCVs, labeled by LAMP1, were RETREG1-positive (Figure 5C). To further confirm whether RETREG1 contribute to xenophagosomes formation, the numbers of xenophagosomes were counted in WT and *RETREG1* knockout (KO) HeLa cells. The results revealed that the numbers of xenophagosomes indicated by LC3B staining reduced in *RETREG1* KO cells (Figure 5D, E). Moreover, ER marker, REEP5 also showed stronger colocalization with LC3B in WT cells compared to *RETREG1* KO cells (Figure 5D). In *retreg1* KO mouse bone-marrow-derived macrophages (BMDMs), REEP5-surrounded *Salmonella* also significantly reduced compared to WT BMDMs (Figure S4E-G). Since xenophagy machinery usually recognizes cytosolic *Salmonella* for clearance, we further detected bacterial growth. The plating assay showed that the colony-forming units (CFUs) significantly increased in *RETREG1* KO HeLa cells compared to WT cells after *Salmonella* infection (Figure 5F). The imaging analysis revealed the multiplication of cytosolic *Salmonella,* which only expressed green florescence in cytosol, increased in *RETREG1* KO HeLa cells compared to WT cells (Figure 5G,H). These results suggest that host cells use RETREG1-associated ER membrane to promote xenophagosomes formation for bacterial clearance.

**Figure 5.**
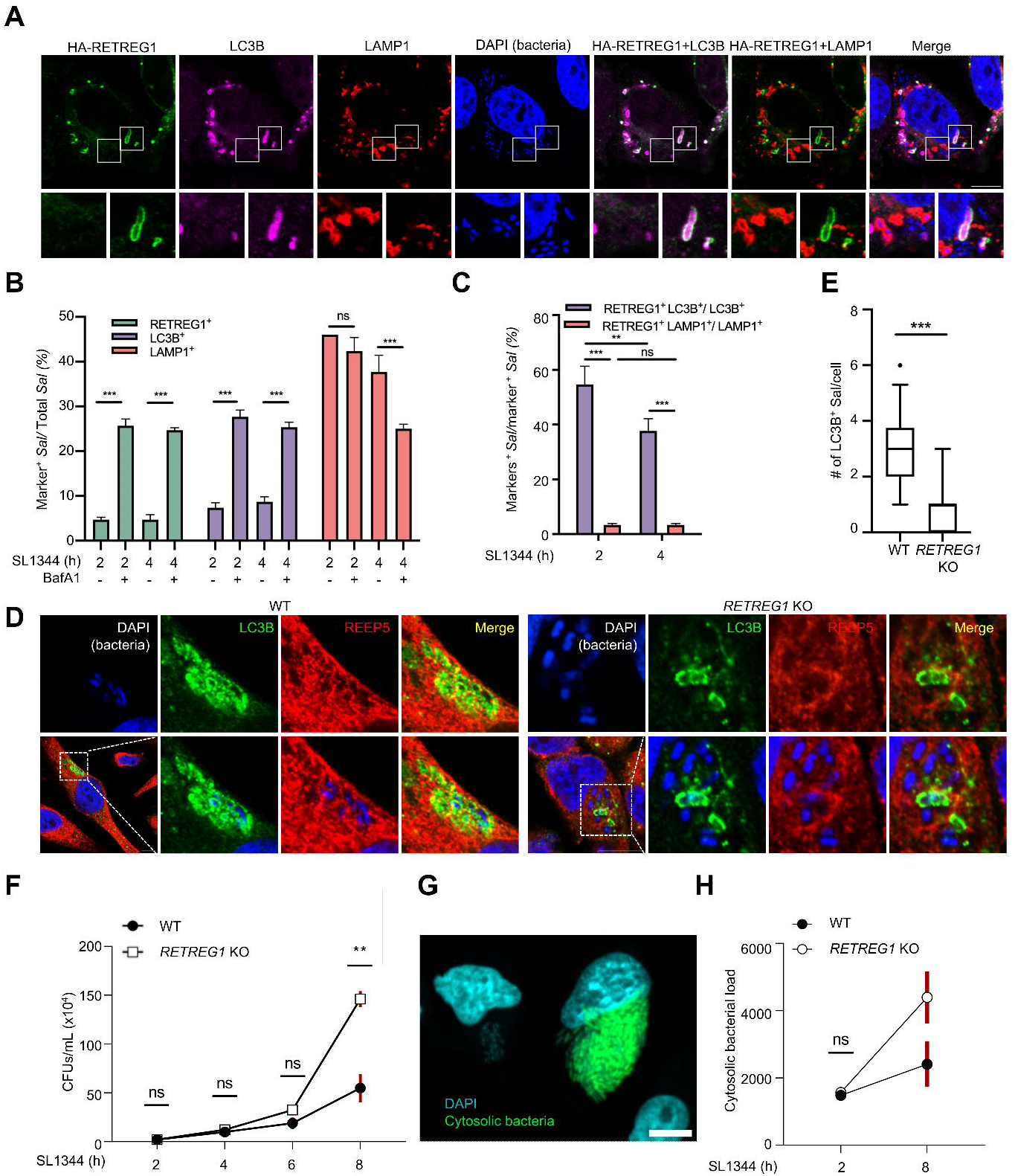
RETREG1 promotes xenophagosomes formation and *Salmonella* clearance. (**A**) Immunofluorescence staining of HA, LC3B, and LAMP1 in HeLa TREx inducible cell line expressing FLAG-HA-RETREG1 infected with SL1344 for 4 h. DAPI was used for cell nuclear and bacterial DNA staining. Scale bar: 10 μm. (**B** and **C**) The percentage of RETREG1^+^, LC3B^+^, or LAMP1^+^ *Salmonella* in total *Salmonella* (**B**), as well as the percentage of RETREG1- and LC3B-double-positive (RETREG1^+^ LC3B^+^) *Salmonella* in all LC3B^+^ *Salmonella* and the percentage of RETREG1- and LAMP1-double-positive RETREG1^+^ LAMP1^+^) *Salmonella* in all LAMP1^+^ *Salmonella* (**C**) was quantified using CellPathfinder software. Data presented as mean + sem, n=3, > 400 cells. Differences were statistically analyzed by one-way ANOVA and Tukey’s multiple comparison test. ***p < 0.001, ns, not significant. (**D** and **E**) Immunofluorescence staining of LC3B and REEP5 in RETREG1 WT HeLa and *RETREG1* KO HeLa infected with SL1344 for 4 h (**D**). DAPI was used for cell nuclear and bacterial DNA staining. Scale bar: 10 μm. Quantification of LC3B^+^ *Salmonella* per cell (**E**). Data were collected 60 cells. Differences were statistically analyzed by unpaired t-test. ***p < 0.001. (**F**) Colony-forming assay was performed to quantify the bacteria count in RETREG1 WT HeLa and *RETREG1* KO HeLa infected with SL1344 100 moi at the indicated time points. Data collected from three independent biological replicates. (**G**) Representative image of HeLa cells infected with pHut-gfp *Salmonella* bacteria after infection (GFP-expressing bacteria are cytosolic, right; vacuolar bacteria have no GFP signal, left). Scale bar 10µm. (**H**) Quantification of GFP-expressing pHut-gfp *Salmonella* bacteria (cytosolic) after infection with moi 100 of RETREG1 WT HeLa and *RETREG1* KO HeLa at the indicated time points. Data collected for >100 cells from three independent biological replicates. (**F**, **H**) Differences were statistically analyzed by two-way ANOVA and Šidák correction for multiple comparisons. **p<0.01, ns not significant.

### OMVs are required for LPS delivery to induce RETREG1-mediated ER remodeling

Transmission electron microscopy (TEM) studies revealed that LPS secreted by *Salmonella* is presented on vesicular structures that resemble OMVs generated by bacteria [20]. OMVs, which are enriched in LPS on the surface, are also responsible for delivery of LPS from extracellular bacteria to the host cell cytosol via endocytosis [25]. Hence, we investigated whether OMV production is required for RETREG1-mediated ER fragmentation. OMV production is governed by various mechanisms that detach the outer membrane from the underlying peptidoglycan and then allow outward budding to form vesicles [27,28]. For example, OmpA (outer membrane protein A) is the major outer membrane porin in *E. coli* and *S. enterica* and is linked to peptidoglycan. Lack of OmpA results in decreased stability of the outer membrane and increased release of OMVs, which has been previously reported for *S. enterica* serovars Typhimurium [37] and Typhi [38], and was confirmed for the mutant strains used in our study by ultrastructural analyses (Figure S5A,B,C). Another example is TolAB, which encodes components of the multi-subunit Tol-Pal complex that controls the constriction of the cell envelope during cell division. Mutant strains lacking TolAB show increased OMV production due to defects in partitioning the outer membrane during cell elongation and septation [39]. DegS is also involved in OMV production. It is part of a sensor system responding to periplasmic stress, which is, for example, induced by accumulation of misfolded secretory proteins. If the stress response system is absent, the accumulation of misfolded proteins in the periplasm induces increased OMV production to remove toxic aggregates [38]. On the contrary, PagL is a deacylase that removes one lipid from hexa- or hepta-acylated lipid A of LPS [40]. This modification alters topology of LPS in the outer membrane, resulting in increased membrane curvature and OMV release [41]. We therefore established *Salmonella* strains mutated in the genes *pagL*, *ompA*, *tolAB*, and *degS,* and obtained the images of OMVs production by scanning electron microscopy (SEM) and TEM followed by measurement of OMVs amount and size distribution (Figure S5A, B, C). Compared to the WT strain, the Δ*ompA*, Δ*tolAB*, and Δ*degS* strains showed increased production of OMVs. Conversely, no vesicles could be detected with the Δ*pagL* strain (Figure S5C), which is in line with previous findings showing that deletion of *pagL* results in decreased generation of OMVs by intracellular *Salmonella* [41].

Mutation of *ompA*, *tolAB*, and *degS*, associated with increased OMVs production, augmented the number of RETREG1- and LC3B-double-positive puncta to varying degrees (Figure 6A and S5D). Especially, mutation of *tolAB* strongly increased puncta formation compared to the WT group. Conversely, the number of RETREG1- and LC3B-double-positive puncta decreased when *pagL* was mutated (Figure 6A and S5D). The LPS levels in the cell lysates and the amount of LPS interacting with RETREG1 positively correlated with the number of RETREG1- and LC3B-double-positive puncta (Figure 6A, C). The bacterial counts for the different isogenic mutant strains did not show significant change at 2 h post-infection (Figure 6B). To further explore whether OMVs can directly activate RETREG1-mediated ER fragmentation, we purified OMVs from *Salmonella* and treated HeLa TREx FLAG-HA-RETREG1 cells with these OMVs. After treatment with *Salmonella* OMVs, we observed colocalization of LPS and HA-RETREG1. The number of RETREG1- and LC3B-double-positive puncta also increased (Figure 6D, E). To differentiate whether RETREG1 mainly bound to shed LPS or the LPS on OMVs, we stained OMVs by Vybrant DiO dye after purification and then treated the cells. Confocal microscopy showed that RETREG1 mainly colocalized with the LPS free from the Vybrant DiO-positive OMVs (Figure S6A). Nevertheless, whether RETREG1 interact with the OMVs or the outer membrane of *Salmonella* still need to be further investigated. Especially, we found that the bacterial outer membrane proteins, such as SlyB and OmpA, are identified in the interactome of RETREG1 during *Salmonella* infection (Figure S7A, B, C). The gene ontology analysis of the bacterial proteins identified by RETREG1-interactome also showed highly correlation with bacterial cell outer membrane components (Figure S7D). Furthermore, the O-Ag mutant strains-derived OMVs lost the ability to trigger RETREG1-mediated ER fragmentation (Figure 6F). We further tested whether extracellular bacteria, more specifically Enteropathogenic *Escherichia coli* (EPEC), can induce a similar phenomenon. In addition to direct exposure, transwells with 0.45-μm polycarbonate membranes were used to prevent bacteria passage but allow OMVs transition. The results reveal that EPEC induced similar amount of RETREG1- and LC3B-double-positive puncta in direct infection and transwell system (Figure 6G, H). The *E. coli* strain without O-Ag, BL21, was used as the negative control, showing no significant formation of RETREG1 puncta (Figure 6G, H). The number of LC3B-decorated RETREG1 puncta also increased in the cells with EPEC-derived OMVs, which were pre-stained by Vybrant DiO after purification. However, BL21-derived OMVs did not cause same effect (Figure 6I and S6B). These results suggest that intracellular and also extracellular bacteria utilize OMVs for the delivery of LPS to activate RETREG1-mediated ER fragmentation.

**Figure 6.**
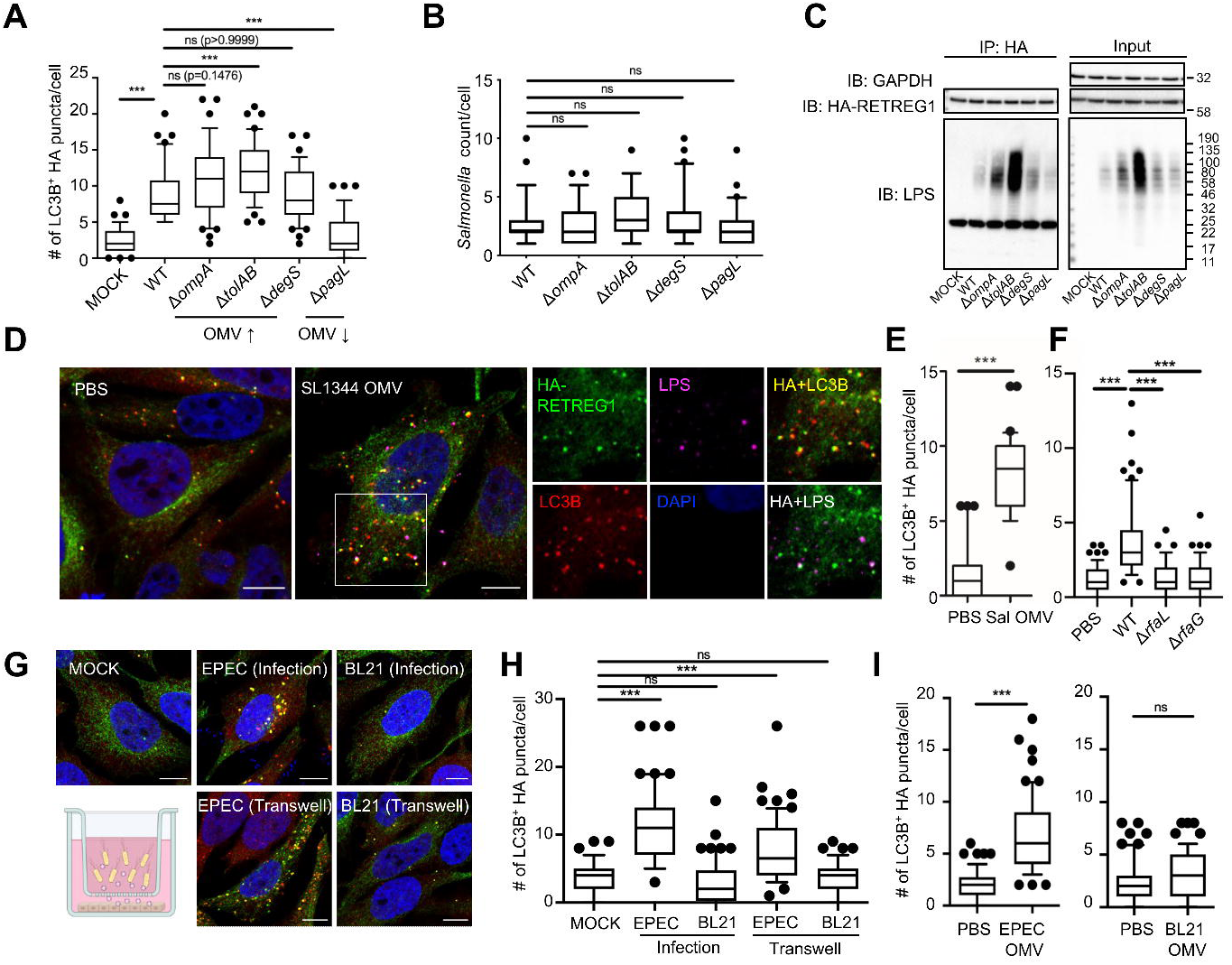
LPS for RETREG1-mediated fragmentation is delivered by OMV. (**A** and **B**) Immunofluorescence staining of HA, *Salmonella* LPS, and LC3B in cells infected with different mutant strains, including Δ*ompA*, Δ*tolAB*, and Δ*degS*, and Δ*pagL* strains. DAPI was used for cell nuclear and bacterial DNA staining. Scale bar: 10 μm. Quantification of LC3B^+^ HA-RETREG1 puncta (**A**) and *Salmonella* count per cell (**B**). Representative images are shown in Figure S5D. (**C**) Co-IP of LPS with HA in cells infected with NCTC12023 WT or indicated mutant strains for 2 h. GAPDH was used as an internal control of input. (**D** and **E**) Immunofluorescence staining of HA, *Salmonella* LPS, and LC3B in cells treated with purified SL1344 OMVs for 6 h (**D**). DAPI was used for cell nuclear DNA staining. Scale bar: 10 μm. Quantification of LC3B^+^ HA-RETREG1 puncta per cell (**E**). (**F**) Immunofluorescence staining of HA and LC3B in cells treated with purified OMVs form NCTC12023 WT or indicated mutant strains for 6 h. DAPI was used for cell nuclear DNA staining. Scale bar: 10 μm. Quantification of LC3B^+^ HA-RETREG1 puncta per cell. (**G** and **H**) Immunofluorescence staining of HA and LC3B in cells infected by EPEC or BL21 directly or with 0.45 μm transwell for 4 h (**G**). DAPI was used for cell nuclear DNA staining. Scale bar: 10 μm. Quantification of LC3B^+^ HA-RETREG1 puncta per cell (**H**). (**I**) Immunofluorescence staining of HA and LC3B in cells treated with purified OMVs form EPEC or BL21 for 6 h. Quantification of LC3B^+^ HA-RETREG1 puncta per cell with OMVs, stained by DiO Vybrant. Representative images are shown in **Figure S6B**. HeLa TREx inducible cell lines expressing FLAG-HA-RETREG1 was used for all experiments in Figure 6. (**A**, **B**, **E**, **F**, **H**, and **I**) Solid bars of boxes indicate the medians. Boxes represent interquartile range from 25^th^ to 75^th^ percentile, and whiskers indicates 10^th^ to 90^th^ percentile. Differences were statistically analyzed by one-way ANOVA and Tukey’s multiple comparison test. ***p < 0.001, ns, not significant. Data were collected 60 cells from two (**A**, **B** and **H**) three independent (**E**, **F** and **I**) biological replicates.

**Figure 7.**
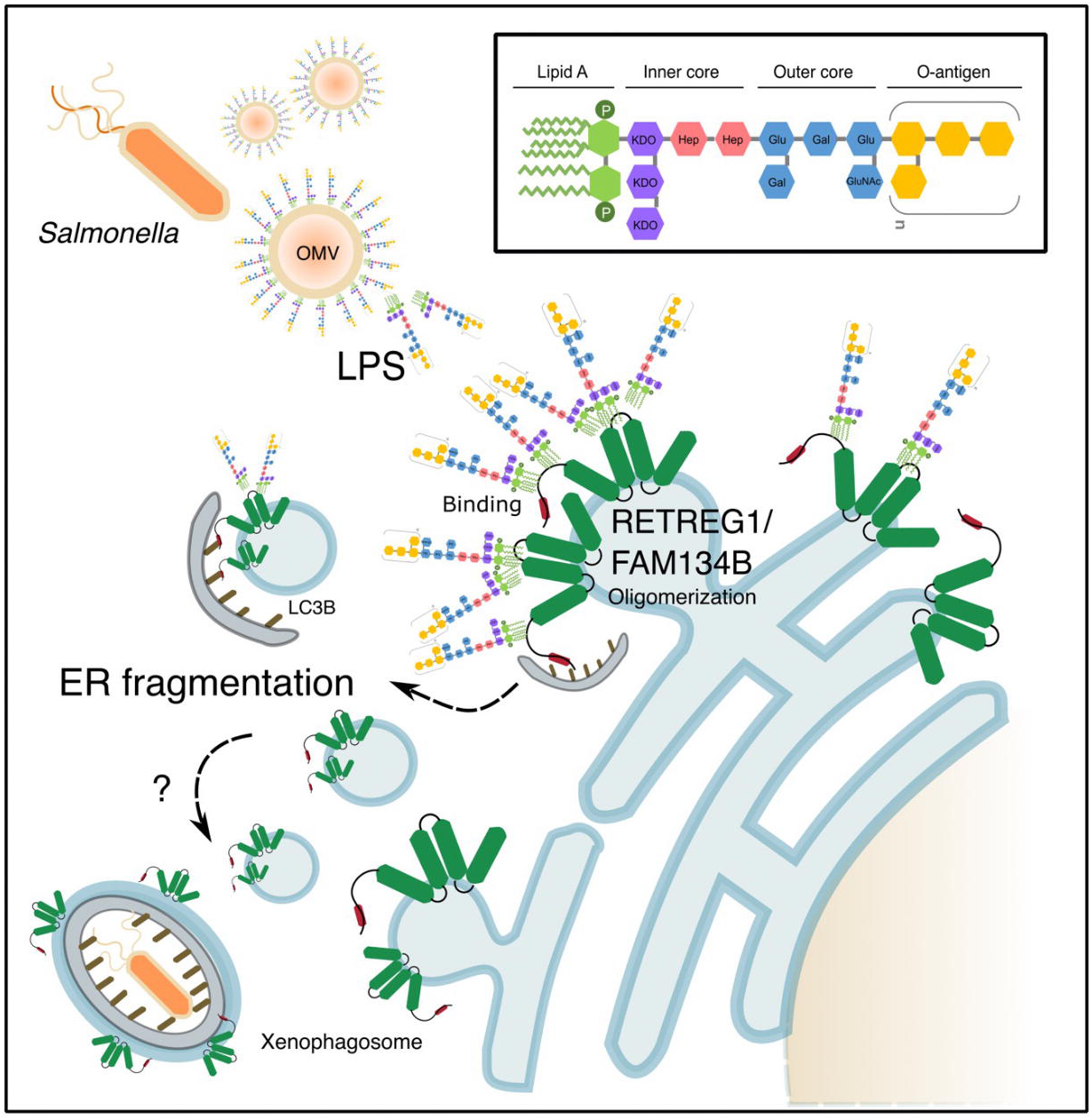
A hypothetical model. Cytosolic LPS binds to the ERphagy receptor RETREG1, triggering oligomerization of RETREG1 and ER fragmentation, and subsequently, the formation of RETREG1- and ER membrane-positive xenophagosome for bacterial clearance. The figure was created by Inkscape.

## Discussion

In this study, we demonstrate that RETREG1, an ERphagy receptor, directly interacts with LPS to induce ER membrane fragmentation and promote the formation of ER membrane-associated xenophagosomes. This interaction is predominantly mediated by lipid A moiety of LPS and positively charged amino acid residues within RETREG1. Additionally, O-Ag component of LPS appears to be critical for oligomerization of RETREG1 and the subsequent membrane fragmentation, although the precise molecular dynamics of this process remain to be elucidated.

Previously we reported that the RHD of RETREG1 forms wedge-shaped membrane inclusions that induce membrane curvature followed by membrane fragmentation [15]. In our recent study [18] We further showed that ubiquitination of the RHD fine-tunes this membrane remodeling activity, enhancing large-scale membrane deformation, increasing receptor clustering, and promoting ERphagy flux. Here, we find that mutations of multiple lysine residues of RETREG1, previously identified as ubiquitination sites, significantly impair LPS binding (Figure 2). This suggest that LPS may functionally utilize ubiquitination in facilitating RETREG1 oligomerization and membrane fragmentation.

The ability of LPS to directly engage RETREG1 and induce ERphagy supports a model wherein *Salmonella* infection activates ERphagy to supply membranes for xenophagosome formation. Our knockdown experiments indicate that depletion of RETREG1 reduces xenophagosome formation, potentially allowing bacterial multiplication. Recently, Gatica *et al.* also showed that *Salmonella* inhibits starvation-induced ERphagy by affecting RETREG1 oligomerization, and that RETREG1 controls *Salmonella* infection through unknown mechanisms [42]. Their study, performed in starved HEK293T cells with elevated baseline ERphagy, identified the *Salmonella* effector SopF as a negative regulator of RETREG1 oligomerization via ADP-ribosylation. In contrast, we did not detect SopF among the RETREG1 interactors in our system (Figure S7A–C), suggesting that *Salmonella* may differentially deploy effectors depending on the host cell type or its nutritional and autophagic status. Indeed, CFU assays revealed a ten-fold higher bacterial load in HEK293T cells compared to HeLa cells. The discrepancy may reflect the opposing modes of RETREG1 regulation, suppression in HEK293T versus activation in HeLa cells, underlying distinct infection outcomes [42]. Future studies should explore whether SopF interferes with LPS-induced ERphagy directly.

Although both our study and that of Gatica et al. demonstrate that RETREG1 contributes to *Salmonella* clearance in certain cell models as well as in *retreg1* KO mice, whether the level for bacterial clearance is sufficient to influence overall pathogenesis upon infection still need to be further investigated [42]. Beyond its role in vacuole formation for bacterial clearance, this ER remodeling pathway likely serves additional functions. The ER is a central hub for various physiological processes, including vesicular trafficking, the unfolded protein response/UPR, ER-lysosome associated degradation/ERAD, and ER-organelle contact-site formation [2]. Dynamics reshaping of the ER enables cells to adopt to stress conditions, for instance, Gram-positive bacterial infections trigger stress via activation of the innate sensor STING1 (stimulator of interferon response cGAMP interactor 1), while simultaneously inducing autophagy [43]. Upon these conditions, ER proteins co-fractionate with LC3B, suggesting activation of ERphagy as a homeostastic response [43]. Whether RETREG1-mediated ER remodeling contributes to analogous responses during Gram-negative bacterial infection remains an intriguing question for future investigation.

Our immunofluorescence experiments showed that the RETREG1 puncta predominantely colocalized with LPS that was not associated with OMVs (Figure S6A). While the precise nature of this free LPS remains unclear, insights can be drawn from studies on GBP1 (guanylate binding protein 1). GBP1 binds directly to lipid A in LPS micelles or on bacterial surfaces and undergoes O-Ag-dependent polymerization [44,45]. Analogously, we observed that RETREG1 binds lipid A moieties and requires O-Ag for oligomerization. Moreover, RETREG1 localized not only to secreted LPS, likely present in micellar form, but also to the bacterial surface, thus mirroring GBP1 behavior. We also detected LPS puncta at early infection stages, before substantial bacterial accumulation, raising questions about the source and mechanism of early LPS release. Several studies suggest that GBPs, along with IRGB10/Gm12250, are instrumental in LPS release into the host cytosol. GBPs are recruited to intracellular pathogens and disrupt pathogen-containing vacuoles, releasing bacteria and their ligands into the cytosol [46]. Subsequently, GBPs recruit IRGB10 to the bacterial surface, disrupting the bacterial cell membrane and releasing bacterial ligands, including LPS [47]. Although we observed a correlation between OMV production by *Salmonella* mutants and intracellular LPS levels, the precise mechanisms governing OMV-mediated LPS release inside host cells remain to be clarified.

In summary, we identify RETREG1 as a novel LPS-binding protein that undergoes oligomerization upon LPS engagement, thereby triggering ER fragmentation and promoting the formation of ER-derived xenophagosomes. Our findings uncover a previously unrecognized host cell response, in which the ER undergoes remodeling by direct interaction of LPS with ERphagy receptor during Gram-negative bacterial infection. This work opens new avenues for understanding how host membrane dynamics regulated in response to bacterial infection and highlights RETREG1 as a potential target for modulating ERphagy during infection.

## Materials and Methods

### Mammalian cell culture

HeLa and HEK293T cells were obtained from American Type Culture Collection (CCL-2, CRL-3216, respectively). WT and *retreg1* KO MEFs and BMDMs were isolated from respective mice genotypes. Animal experiments are approved by the Thüringer Landesamt für Lebensmittelsicherheit und Verbraucherschutz (TLV) (license number UKJ 22-019). MEFs were isolated and cultured from E13.5 stage embryos and SV40 large T antigen plasmid was used to immortalize MEFs. To culture BMDMs, femur and tibia bones were isolated from 12–16-week-old WT and *retreg1* KO mice after euthanasia. The bones were cleaned, sterilized, and the bone marrow was flushed out with sterile PBS (Gibco, 14190094). It was centrifuged at 300 x g for 5 min and resuspended with BMDM culture media (DMEM supplemented with 10% fetal bovine serum, 1% penicillin-streptomycin and 20 ng/mL of murine CSF1 (colony stimulating factor 1 (macrophage)) and plated in Petri dishes. Fresh media was added on day 2. By day 5, differentiated macrophages were attached and the debris were removed, the cells were washed with 1x PBS, collected, counted, and plated for the experiments. Flp-In HeLa TREx cell lines (Invitrogen, R71407) were used to generate inducible cell lines by cloning the cDNAs of *RETREG1*, *RETREG1* mtLIR (DDFELLD LIR substituted with seven alanine residues), and RTN3L into MSCV iTAP N-FLAG-HA retroviral vector (Addgene, 41033; deposited by Wade Harper). Then, 2 μg/ml puromycin and 15 μg/ml blasticidin were used for selection and maintenance of stable cells. Cells were cultured in Dulbecco’s Modified Eagle’s Medium (DMEM; Gibco, 10564011) supplemented with 10% fetal bovine serum, 1% penicillin-streptomycin at 37°C with 5% CO_2_. BMDMs were additionally supplemented with 20 ng/mL of murine CSF1 (Miltenyi BioTech, 130-101-706).

### Microbe strains

*S.* Typhimurium strains used in this study. The following isogenic mutant and complemented NCTC12023 strains were established by Prof. Dr. Michael Hensel (Universität Osnabrück, Osnabrück, Germany) [31].

1. SL1344 (WT)
2. NCTC12023 (WT)
3. NCTC12023 strains with deletion of *ompA, tolAB, degS* or *pagL*. Mutant strains of *S. enterica* serovar Typhimurium strain NCTC12023 were generated by Red-mediated homologous recombination basically as described previously [48], using pWRG730, developed by Hoffmann et al, was used as helper plasmid for controlled expression of Red functions [49]. The 60-mer oligonucleotide sets as listed in Table S1 were used to amplify the *aph* targeting cassette using pKD13 (for *tolA*, *degS*, or *ompA*; ECGRC, CGSC: 7633) or pKD4 (for *pagL*; ECGRC, CGSC: 7632) as template DNA. Proper insertion of *aph* cassettes into target genes was confirmed by colony PCR using gene-specific check primers flanking deletions, and universal K1 Red Del. Confirmed mutated alleles were moved into fresh strain background using P22 transduction. The effects of mutations on outer membrane integrity were analysed by SEM and nsTEM of strains grown overnight on LB agar (Fig. S5A, B)
4. NCTC12023 strains with deletion on *rfbP, rfaL, rfaG,* or *rfaH*. Mutant of *rfbP* (O-Ag transferase; NP_461027; Δ*rfbP*) and *rfaL* (O-Ag ligase; NP_462613; Δ*rfaL*) are defective in the addition of the O-Ag to the outer core. Mutant of *rfaG* (glucosyltransferase I; NP_462622; Δ*rfaG*) caused outer core cannot be added to inner core. Mutant of *rfaH* (transcriptional activator; NP_ 462862; Δ*rfaH*), which is the gene regulates the stability of the long mRNA for LPS biosynthesis genes, also reduced polymeric O-Ag.
5. NCTC12023 strains with complementation of deletion of *rfaL* and *rfaG*
6. pHut-gfp *S.* Typhimurium SL1344 bacteria (express GFP when cytosolic)

*E. coli* strains used in this study.

1. NEB Turbo (New England Biolabs, C2984I)
2. EPEC O127:H6 (A kind gift from Prof. Dr. Volkhard A. J. Kempf [Goethe University Frankfurt, Frankfurt, Germany])
3. BL21 (DE3) (New England Biolabs, C2527H)

Chemically competent NEB Turbo cells were used for plasmid amplification.

### Bacterial infection

*S.* Typhimurium and *E. coli* were cultured overnight and then diluted at ratio of 1 to 33 in fresh LB broth. The diluted cultures were then grown at 37°C for 3 h. For infection, the resuspended bacteria were adjusted to OD600 of 1 and added to HeLa cells or MEFs at a multiplicity of infection (MOI) of 100, and BMDMs at a MOI of 10 without antibiotics. After 30 min of infection, media were replaced by medium with 100 μg/ml gentamicin to remove extracellular bacteria. Samples were collected at the indicated time points after infection. For colony forming assay, which was performed as previously described [50], after 10 min of *S.* Typhimurium infection, the cells were washed with PBS twice to removed extracellular bacteria and incubated in antibiotics-free DMEM for 20 min. Then, the media were changed to DMEM containing 50 μg/ml gentamicin for 40 min and replaced by DMEM containing 5 μg/ml gentamicin. The cells were lysed with lysis buffer (1% Triton X-100 [Sigma-Aldrich, X100] and 0.1% SDS in PBS) at indicated time points and serial diluted with PBS and plated onto LB plates. After 24-h incubation at 37°C, the colonies were calculated.

### Immunofluorescence staining and confocal imaging

Cells which seeded in the 6-well plates with cover slips fixed by 4% paraformaldehyde in PBS for 15 min at room temperature. The buffer with 0.1% saponin (Sigma-Aldrich, SAE0073-10G), 1% bovine serum albumin (BSA; Carl Roth, 8076.3), 0.05% NaN3 in PBS was used for permeabilization and blocking for 1 h at room temperature. Samples were then probed with the indicated primary antibodies as well as Alexa Fluor 488, Alexa Fluor 546, and Alexa Fluor 633 fluorescently labelled secondary antibodies (Invitrogen, A-11008, A-21052, A-11077). After DAPI staining for nucleus and bacteria DNA, the samples were fixed by mounting medium. Confocal imaging was done using a Zeiss LSM 780 confocal microscope and analyzed by Zeiss Zen (black edition) software. For high-throughput confocal imaging, cells were seeded in the 96-well black plates with glass bottom followed by immunofluorescence staining and imaging by CellVoyager CQ1 (Yokogawa) system and quantified using CellPathfinder software (version 3.06.01.08).

### Immunoprecipitation and western blotting

For immunoprecipitation, cells were lysed in the lysis buffer containing 50 mM Tris-HCl, pH 7.5,150 mM NaCl, 5 mM EDTA, and 1% NP-40 (Sigma-Aldrich, NP-40) with protease inhibitor cocktail (Roche, 11836170001). Lysates were incubated with 15 μl anti-HA agarose beads (Santa Cruz Biotechnology, sc-500777) overnight at 4°C with end-to-end rotation. Beads were washed three times with wash buffer containing 50 mM Tris-HCl, pH 7.5, 400 mM NaCl, 5 mM EDTA). Proteins were eluted with 2X loading dye by boiling for 10 min followed by western blotting. For western blotting, cells were lysed with lysis buffer containing 50 mM Tris-HCl, pH 7.5, 150 mM NaCl, 5 mM EDTA, and 1% Triton X-100 with protease inhibitor cocktail (Roche, 11836170001). After protein quantification, samples were boiled with loading dyes for 10 min. Samples were separated by running SDS-PAGE with Tris-Glycine gels (Mini-PROTEAN TGX Stain-Free Precast Gels; Bio-Rad, 4561096) followed by transfer to PVDF membranes, which were then blocked with 5% milk in TBST (TBS [Bio-Rad, 1706435] with 0.1% Tween 20 [Sigma-Aldrich, 11332465001]). After blocking, the membranes were sequentially hybridized with primary and HRP (horseradish peroxidase)-conjugated secondary antibodies (anti-rabbit-HRP [Dako, P0448], mIgGk BP-HRP [Santa Cruz Biotechnology, sc-516102], anti-rat-HRP [Cell Signaling Technologies, 7077S]). The chemiluminescence method was performed to detect proteins expression.

### Transfection of siRNA

Cells were seeded in the 6-well plates on a day before experiments. *RETREG1* siRNA and Scramble siRNA (Eurofins) were transfection using lipofectamine RNAiMAX (Invitrogen, 13778100) as manufacturer’s instruction. After 48-h culture, cells were reseeded for the following experiments.

### Streptavidin affinity-isolation assay

To immobilize LPS onto streptavidin agarose beads, the indicated amount of biotin-conjugated *E. coli* O111:B4 LPS (InvivoGen, tlrl-lpsbiot) were added to 10 μl of streptavidin agarose beads (Cytiva, 45-000-279) and incubated at 4°C with rotation for 1 h. The beads were then washed 3 times with the lysis buffer containing 50 mM Tris-HCl, pH 7.5, 150 mM NaCl, 5 mM EDTA, and 1% NP-40 to remove unconjugated ligands. Inducible TREx HeLa cells of FLAG-HA-RETREG1 and FLAG-HA-RTN3 induced by treatment of 1 μg/ml doxycycline (Thermo Fisher Scientific, J60422.06) for 24 h or HEK cells transfected with the indicated plasmids were lysed and collected in the lysis buffer supplemented with a protease inhibitor mixture. Cell lysates were precleared by incubating with streptavidin Sepharose (Cytiva, 45-000-279) at 4°C with rotation for 1 h. Supernatants were transferred to a new microtube and incubated with biotin-LPS-immobilized streptavidin agarose beads (Cytiva, 45-000-279) overnight at 4°C with rotation. The beads were then washed three times with wash buffer containing 50 mM Tris-HCl, pH 7.5, 400 mM NaCl, 5 mM EDTA and eluted in 2X loading dye by boiling for 10 min followed by western blotting to detect the binding proteins. For competition assay, 50 μg of unlabeled S-LPS, Ra-LPS, Re-LPS, lipid A, and MDP were first incubated with pre-cleared lysates at 4°C for 1 h with rotation before adding biotinylated LPS-immobilized streptavidin beads. For examination of polymyxin B effect, the indicated concentrations of polymyxin B were first added to biotinylated LPS-immobilized streptavidin beads and incubate with rotation for 0.5 h. After twice washes by lysis buffer, the pre-cleared lysates were added.

### RETREG1 protein purification

For purification of RETREG1 protein from HEK293T cells, the plasmid of S protein-FLAG-Streptavidin binding peptide (SFB; Addgene, 99391; deposited by Michael Huen)-RETREG1 was transfected into HEK293T cells. Cells were lysed with the lysis buffer containing 50 mM Tris-HCl, pH 7.4,150 mM NaCl, 0.5 mM EDTA, and 1% Triton X-100. After centrifugation at 17,000 x g, 15 min, the lysates were collected and incubated with streptavidin Sepharose at constant agitation at 4°C, overnight. After washing three time with the buffers (50 mM Tris, pH=7.4, 150 mM NaCl, 0.5 mM EDTA) containing 0.5% NP-40, 0.3% NP-40 and 0.1% NP-40, respectively, the buffers were removed and the SFB-RETREG1 protein on the Sepharose was further eluted by incubated with 2 mM D-biotin (Thermo Fisher Scientific, B20656) dissolved in the buffer containing 50 mM Tris-HCl, pH 7.4,150 mM NaCl, 0.5 mM EDTA, and 0.1% NP-40 for 1 h at 4°C with constant agitation at 800 rpm. The eluted SFB-RETREG1 was collected for blue native page analysis.

Recombinant GST-RETREG1 was purified from *E. coli* for Liposome-remodeling assay as previous described [18]. Briefly, RETREG1-GST plasmids were transformed into BL21DE3 strain of *E. coli*, and a single colony was used to grow the primary culture in the presence of appropriate antibiotic (ampicillin). The overnight grown primary cultures were used to inoculate secondary cultures and allowed to grow up to an OD of 0.6 by shaking at 200 rpm and 37°C, followed by induction of the cultures using 0.5 mM of IPTG (Santa Cruz Biotechnology, sc-202185B) for 3 h. The cells were harvested, suspended in ice cold PBS and lysed using sonication, centrifuged (10,000 x g for 10 min) to separate the supernatant and pellet. The resultant supernatant was further fractioned by ultracentrifugation at 80,000 x g for 100 min at 4°C. The pellet fractions were dissolved in PBS buffer containing dodecyl β-D-maltoside (DDM, 0.05%; Anatrace, D310), followed by affinity purification using glutathione-Sepharose TM4 Fast Flow columns (GE Healthcare, GE17-5132-01). The protein was eluted from the column using PBS containing 10 mM reduced glutathione and 0.05% DDM. The fractions containing RETREG1-GST protein were concentrated and buffered exchanged with storage buffer (50 mM HEPES, pH 7.5, 150 mM NaCl) containing 0.0075% DDM.

### Blue native page

Blue native page analysis was performed as previously described with slight modifications [17,30]. SFB-RETREG1 protein was incubated with indicated amount of S-LPS or lipid A at 37°C for 2 h. Then, the mixture was incubated with 2 μl 50% glycerol and 3 μl 0.1% Coomassie Brilliant Blue G-250 for 15 min at room temperature followed by loading to 4%–12% gradient native PAGE gel (Bio-Rad, 3450123), running in blue cathode/anode buffer (Invitrogen, BN2007) for 2 h at 150 V in an ice bath. The gels were further transfer to PVDF membranes by electroblotting followed by detection with anti-FLAG antibody.

### Liposome preparation

Liposomes were prepared as previously described [15]. In brief, two kinds of lipids,1,2-dioleoyl-sn-glycero-3-phosphocholine (DOPC; Avanti Polar Lipids Inc., 850375) and 1,2-dioleoyl-sn-glycero-3-phosphoethanolamine (DOPE; Avanti Polar Lipids Inc., 850725), were dissolved in a mixture of chloroform and methanol (4:1) in a glass flask to obtain the molar ratio of 0.8:0.2 (DOPC:DOPE). The organic solvent was then evaporated to get a dry lipid film, which was then hydrated with liposome buffer (50 mM HEPES, 150 mM NaCl buffer at pH 7.4) to a final concentration 15 mg/ml lipid solution. The liposomes were formed by sonication in an ultrasound bath followed by flash-freeze in liquid nitrogen and slowly thaw at room temperature for 10 cycles. The hydrated liposomes were further extruded by a lipid extruder (Avanti Polar Lipids Inc.) with 200 nm polycarbonate membranes (Avanti Polar Lipids) for 21 times.

### Liposome-remodeling assay

A liposome-remodeling assay was performed as previous described with minor modifications [15]. Liposomes and the purified RETREG1-GST protein were mixed at an indicated lipid-to-protein ratio (LPR) and incubated for 18 h at 22°C in 30 μL of liposome buffer with constant agitation (600 rpm). To investigate the effect of different LPS, RETREG1-GST protein was first added to liposome at LPR 100:1 for 1 h, then added LPS to the proteoliposome and incubated for 17 h at 22°C. All the samples were assessed by negative-stain transmission electron microscopy. For imaging, samples were first diluted to a final lipid concentration of 1 mg/ml by liposome buffer. Five μl of each diluted sample was deposited to the carbon-coated copper grids (SPI Supplies, 3405C-CF), which were already glow-discharged for 20 s at 15 mA and 0.38 mbar vacuum. After 1-min incubation and washes with buffer for twice, 1% uranyl formate was subsequently used to stain the grids for 10 s at room temperature. A 120 kV Tecnai Spirit Biotwin electron microscope (FEI) equipped with a 4k × 4k CCD detector (US4000-1, Gatan) was used to taken 10 micrographs for each sample. Image J software was used to measure the sizes (diameters) of 300 proteoliposomes for each sample.

### OMV isolation

0.5 L overnight grown bacterial cultures were harvested for OMV isolation. Bacterial cells were first removed by centrifugation (12,000 x g for 15 min), the supernatant was filtered through a 0.45-µm PVDF membrane, the filtered broth concentrated to 50 ml (Vivaflow 200, 100,000 MWCO PES; Sartorius, VF20P4), and the OMVs were pelleted by ultracentrifugation at 60,000 rpm (369548.3 x g) for 1 h at 4°C. The OMVs were washed with PBS and pelleted again by ultracentrifugation. The OMVs were resuspended in 0.5 ml PBS. In the cases which needed to do immunofluorescence labeling, Vybrant DiO cell labeling dye (Thermo Fisher Scientific, V22886) was added at 1:50 and incubated at 37°C for 20 min. After labeling, the excess dye was removed and washed with PBS three times at 3000 rpm (956 x g) by Micron 10 kDa centrifugal filter units (Millipore, MRCPRT010) in a benchtop centrifuge. The OMVs were further sterilized by syringe filters (0.22 µm) and measured concentration by bicinchoninic acid assay (BCA) method. OMVs were added to the cells at concentration of 25 µg/ml.

### Quantification and statistical analysis

GraphPad Prism 8 was used to do statistical analysis. Comparisons of two variables were analyzed by unpaired and two-tailed Student’s t tests. For three or more sets of data, statistics were performed by one-way ANOVA with Tukey’s multiple comparison post test. * indicates p < 0.05, ** indicates p < 0.01, *** indicates p < 0.001. N represents the number of independent biological replicates. Data are represented as mean ± sem.

## Acknowledgements

We thank Prof. Dr. Volkhard A. J. Kempf at the Institute for Medical Microbiology and Infection Control, Hospital of the Goethe University for the kind gift of the EPEC O127:H6 strain. We thank the staff at the Quantitative Proteomics Unit at Goethe University for support with mass spectrometry and the Frankfurt Center for Advanced Microscopy (FCAM) for access to microscopes.

## Data availability

The MS proteomics data have been deposited to the ProteomeXchange Consortium through the PRIDE partner repository with the dataset identifiers PXD053709.

## Declaration of generative AI and AI-assisted technologies in the writing process

During the preparation of this work the author(s) used chatGPT in order to improve language and readability for some sentences. After using this tool, the authors reviewed and edited the content as needed and takes full responsibility for the content of the publication.

## Abbreviations

AH: amphipathic helix
BMDMs: bone-marrow-derived macrophages
Co-IP: co-immunoprecipitation
BafA1: bafilomycin A_1_
Cterm: C-terminal region (Cterm)
CFU: colony-forming units
DAPI: 4’,6-diamidino-2-phenylindole
ER: endoplasmic reticulum
EPEC: enteropathogenic *Escherichia coli*
GBP: guanylate binding protein
Gm12250/IRGB10: predicted gene 12250
KDO: keto-3-deoxy-octonate
LPR: lipid-to-protein ratio
LPS: lipopolysaccharide
MAP1LC3B/LC3B: microtubule associated protein 1 light chain 3 beta
mtLIR: LC3B-interacting region mutant
MDP: muramyl dipeptide
OMVs: outer membrane vesicles
O-Ag: O-antigen
OmpA: outer membrane protein A
RHD: reticulum homology domain
R-LPS: rough-LPS
S-LPS: smooth-LPS
SCVs: *Salmonella*-containing vacuoles
SFB: S-protein-FLAG-streptavidin binding peptide
TM: transmembrane domain
TEM: transmission electron microscopy
WT: wild-type.

## References

[1] Escoll P, Mondino S, Rolando M, et al. Targeting of host organelles by pathogenic bacteria: a sophisticated subversion strategy. Nat Rev Microbiol. 2016;14(1):5–19.

[2] Gubas A, Dikic I. ER remodeling via ER-phagy. Mol Cell. 2022;82(8):1492–1500.

[3] Chino H, Mizushima N. ER-phagy: quality control and turnover of endoplasmic reticulum. Trends Cell Biol. 2020;30(5):384–398.

[4] Chino H, Mizushima N. ER-phagy: quality and quantity control of the endoplasmic reticulum by autophagy. Cold Spring Harb Perspect Biol. 2023;15(1).

[5] Khaminets A, Heinrich T, Mari M, et al. Regulation of endoplasmic reticulum turnover by selective autophagy. Nature. 2015;522(7556):354–8.

[6] Kumar D, Lak B, Suntio T, et al. RTN4B interacting protein FAM134C promotes ER membrane curvature and has a functional role in autophagy. Mol Biol Cell. 2021;32(12):1158–1170.

[7] Reggio A, Buonomo V, Berkane R, et al. Role of FAM134 paralogues in endoplasmic reticulum remodeling, ER-phagy, and collagen quality control. EMBO Rep. 2021;22(9):e52289.

[8] Grumati P, Morozzi G, Holper S, et al. Full length RTN3 regulates turnover of tubular endoplasmic reticulum via selective autophagy. Elife. 2017;6.

[9] Fumagalli F, Noack J, Bergmann TJ, et al. Translocon component Sec62 acts in endoplasmic reticulum turnover during stress recovery. Nat Cell Biol. 2016;18(11):1173–1184.

[10] Smith MD, Harley ME, Kemp AJ, et al. CCPG1 Is a non-canonical autophagy cargo receptor essential for ER-phagy and pancreatic ER proteostasis. Dev Cell. 2018;44(2):217–232 e11.

[11] Chen Q, Xiao Y, Chai P, et al. ATL3 Is a tubular ER-phagy receptor for GABARAP-mediated selective autophagy. Curr Biol. 2019;29(5):846–855 e6.

[12] An H, Ordureau A, Paulo JA, et al. TEX264 Is an endoplasmic reticulum-resident ATG8-interacting protein critical for ER remodeling during nutrient stress. Mol Cell. 2019;74(5):891–908 e10.

[13] Chino H, Hatta T, Natsume T, et al. Intrinsically disordered protein TEX264 mediates ER-phagy. Mol Cell. 2019;74(5):909–921 e6.

[14] Nthiga TM, Kumar Shrestha B, Sjottem E, et al. CALCOCO1 acts with VAMP-associated proteins to mediate ER-phagy. EMBO J. 2020;39(15):e103649.

[15] Bhaskara RM, Grumati P, Garcia-Pardo J, et al. Curvature induction and membrane remodeling by FAM134B reticulon homology domain assist selective ER-phagy. Nat Commun. 2019;10(1):2370.

[16] Siggel M, Bhaskara RM, Moesser MK, et al. FAM134B-RHD protein clustering drives spontaneous budding of asymmetric membranes. J Phys Chem Lett. 2021;12(7):1926–1931.

[17] Jiang X, Wang X, Ding X, et al. FAM134B oligomerization drives endoplasmic reticulum membrane scission for ER-phagy. EMBO J. 2020;39(5):e102608.

[18] Gonzalez A, Covarrubias-Pinto A, Bhaskara RM, et al. Ubiquitination regulates ER-phagy and remodelling of endoplasmic reticulum. Nature. 2023.

[19] Park BS, Song DH, Kim HM, et al. The structural basis of lipopolysaccharide recognition by the TLR4-MD-2 complex. Nature. 2009;458(7242):1191–5.

[20] Garcia-del Portillo F, Stein MA, Finlay BB. Release of lipopolysaccharide from intracellular compartments containing *Salmonella* typhimurium to vesicles of the host epithelial cell. Infect Immun. 1997;65(1):24–34.

[21] Shi J, Zhao Y, Wang Y, et al. Inflammatory caspases are innate immune receptors for intracellular LPS. Nature. 2014;514(7521):187–92.

[22] Aglietti RA, Estevez A, Gupta A, et al. GsdmD p30 elicited by caspase-11 during pyroptosis forms pores in membranes. Proc Natl Acad Sci U S A. 2016;113(28):7858–63.

[23] Ding J, Wang K, Liu W, et al. Pore-forming activity and structural autoinhibition of the gasdermin family. Nature. 2016;535(7610):111–6.

[24] Santos JC, Boucher D, Schneider LK, et al. Human GBP1 binds LPS to initiate assembly of a caspase-4 activating platform on cytosolic bacteria. Nat Commun. 2020;11(1):3276.

[25] Vanaja SK, Russo AJ, Behl B, et al. Bacterial outer membrane vesicles mediate cytosolic localization of LPS and caspase-11 activation. Cell. 2016;165(5):1106–1119.

[26] Silhavy TJ, Kahne D, Walker S. The bacterial cell envelope. Cold Spring Harb Perspect Biol. 2010;2(5):a000414.

[27] Schwechheimer C, Kuehn MJ. Outer-membrane vesicles from Gram-negative bacteria: biogenesis and functions. Nat Rev Microbiol. 2015;13(10):605–19.

[28] Juodeikis R, Carding SR. Outer membrane vesicles: biogenesis, functions, and issues. Microbiol Mol Biol Rev. 2022;86(4):e0003222.

[29] Adams PG, Lamoureux L, Swingle KL, et al. Lipopolysaccharide-induced dynamic lipid membrane reorganization: tubules, perforations, and stacks. Biophys J. 2014;106(11):2395–407.

[30] Wittig I, Braun HP, Schagger H. Blue native PAGE. Nat Protoc. 2006;1(1):418–28.

[31] Zenk SF, Jantsch J, Hensel M. Role of *Salmonella enterica* lipopolysaccharide in activation of dendritic cell functions and bacterial containment. J Immunol. 2009;183(4):2697–707.

[32] Madan R, Rastogi R, Parashuraman S, et al. *Salmonella* acquires lysosome-associated membrane protein 1 (LAMP1) on phagosomes from Golgi via SipC protein-mediated recruitment of host Syntaxin6. J Biol Chem. 2012;287(8):5574–87.

[33] LaRock DL, Chaudhary A, Miller SI. *Salmonellae* interactions with host processes. Nat Rev Microbiol. 2015;13(4):191–205.

[34] Wild P, Farhan H, McEwan DG, et al. Phosphorylation of the autophagy receptor optineurin restricts *Salmonella* growth. Science. 2011;333(6039):228–33.

[35] Ravenhill BJ, Boyle KB, von Muhlinen N, et al. The cargo receptor NDP52 initiates selective autophagy by recruiting the ULK complex to cytosol-invading bacteria. Mol Cell. 2019;74(2):320–329 e6.

[36] Perrin AJ, Jiang X, Birmingham CL, et al. Recognition of bacteria in the cytosol of mammalian cells by the ubiquitin system. Curr Biol. 2004;14(9):806–11.

[37] Deatherage BL, Lara JC, Bergsbaken T, et al. Biogenesis of bacterial membrane vesicles. Mol Microbiol. 2009;72(6):1395–407.

[38] Nevermann J, Silva A, Otero C, et al. Identification of genes involved in biogenesis of outer membrane vesicles (OMVs) in *Salmonella enterica* Serovar Typhi. Front Microbiol. 2019;10:104.

[39] Szczepaniak J, Press C, Kleanthous C. The multifarious roles of Tol-Pal in Gram-negative bacteria. FEMS Microbiol Rev. 2020;44(4):490–506.

[40] Kawasaki K, Ernst RK, Miller SI. 3-O-deacylation of lipid A by PagL, a PhoP/PhoQ-regulated deacylase of *Salmonella* typhimurium, modulates signaling through Toll-like receptor 4. J Biol Chem. 2004;279(19):20044–8.

[41] Elhenawy W, Bording-Jorgensen M, Valguarnera E, et al. LPS remodeling triggers formation of outer membrane vesicles in *Salmonella*. mBio. 2016;7(4).

[42] Gatica D, Alsaadi RM, El Hamra R, et al. The ER-phagy receptor FAM134B is targeted by *Salmonella* Typhimurium to promote infection. Nat Commun. 2025;16(1):2923.

[43] Moretti J, Roy S, Bozec D, et al. STING senses microbial viability to orchestrate stress-mediated autophagy of the endoplasmic reticulum. Cell. 2017;171(4):809–823 e13.

[44] Kutsch M, Sistemich L, Lesser CF, et al. Direct binding of polymeric GBP1 to LPS disrupts bacterial cell envelope functions. EMBO J. 2020;39(13):e104926.

[45] Santos NC, Silva AC, Castanho MA, et al. Evaluation of lipopolysaccharide aggregation by light scattering spectroscopy. Chembiochem. 2003;4(1):96–100.

[46] Meunier E, Dick MS, Dreier RF, et al. Caspase-11 activation requires lysis of pathogen-containing vacuoles by IFN-induced GTPases. Nature. 2014;509(7500):366–70.

[47] Man SM, Karki R, Sasai M, et al. IRGB10 liberates bacterial ligands for sensing by the AIM2 and caspase-11-NLRP3 inflammasomes. Cell. 2016;167(2):382–396 e17.

[48] Datsenko KA, Wanner BL. One-step inactivation of chromosomal genes in Escherichia coli K-12 using PCR products. Proc Natl Acad Sci U S A. 2000;97(12):6640–5.

[49] Hoffmann S, Schmidt C, Walter S, et al. Scarless deletion of up to seven methyl-accepting chemotaxis genes with an optimized method highlights key function of CheM in *Salmonella* Typhimurium. PLoS One. 2017;12(2):e0172630.

[50] Steele-Mortimer O. Infection of epithelial cells with *Salmonella* enterica. Methods Mol Biol. 2008;431:201–11.

