## Supplementary information for "Intracellular lipopolysaccharide binds RETREG1/FAM134B to regulate ER remodeling upon bacterial infection"

**METHOD DETAILS**

***Mouse model for S.* Typhimurium *infection***

*Salmonella* *enterica* subsp. enterica serovar Typhimurium ATCC14028 (NCTC12023) carrying a green fluorescent protein (GFP) expression plasmid (pGFP, AmpR, kindly provided by Brendan Cormack, Stanford, USA) were cultured in Luria Bertani (LB) broth overnight at 37°C. The overnight culture was diluted 1∶10 in fresh LB and incubated at 37°C until reaching the logarithmic phase (OD600 approximately 0.5). Bacteria were washed and adjusted to OD600 0.55–0.60 containing approximately 1.5–2.0×10^8^ CFU/ml. One-day-old C57BL/6 animals were infected orally with 10^2^ CFU *S*. Typhimurium in a volume of 1 µl PBS. The administered inoculum was confirmed by serial dilution and plating. At day 4 post-infection, the small intestine tissue was collected and fixed in 4% paraformaldehyde.

***HA-RETREG1 interactome analysis in cells infected with* Salmonella *and sample preparation for MS***

HeLa TREx cells were induced with doxycycline to express HA-FLAG-RETREG1. After 16 h, cells were infected with *Salmonella* for 2 and 4 h. After the infection with *Salmonella*, cells were lysed with 1% Triton X-100 in lysis buffer (50 mM Tris-HCl, pH 7.5, 150 mM NaCl, 0.5 mM EDTA, Roche EDTA-free protease inhibitor cocktail and NEM), followed by incubation with HA-beads (Sigma-Aldrich, 26181) overnight at 4°C on a rotating platform. Protein-bound beads were washed three times with lysis buffer and three times in the same buffer without detergents before the elution. Samples were eluted with 25 µl SDC buffer (2% sodium deoxycholate [Sigma-Aldrich, 264103] and 50 mM Tris-HCl, pH 8.5) and heated to 95°C for 5 min. Additionally, 25 µl of 50 mM Tris-HCl, pH 8.5 containing 1 mM TCEP (Thermo Fisher Scientific, 77720), 4 mM chloroacetamide (Thermo Fisher Scientific, A39270) were added to the samples in order to alkylate and reduce the proteins and incubated to 95°C for 5 min. We then added 500 ng trypsin (Promega, V5280) in 25 µl 50 mM Tris-HCl, pH 8.5 to each sample and incubated overnight at 37 °C. The reaction was stopped with 150 µl of 1% TFA in isopropanol. Peptides were cleaned up using SDB-RPS stage tips (Sigma-Aldrich, CDS Empore, 2241). After one wash with 1% TFA in isopropanol and one wash with 0.2% TFA in water, peptides were eluted in 80% acetonitrile plus 1.25% ammonia. Eluted peptides were dried and processed for LC–MS.

***LC–MS analysis***

Dried peptides were resolved in 2% acetonitrile containing 0.1% TFA and loaded to the 35-cm, 75-µm ID fused-silica column (Thermo Fisher Scientific, ESS903), which packed in house with 1.9-µm C18 particles (Reprosil pur; Dr. Maisch, r119.b9), then analyzed on a QExactive HF mass spectrometer coupled to an easy nLC 1200 (Thermo Fisher Scientific). An integrated column oven (Sonation) was used to maintain the column at 50°C. Peptides were eluted using a non-linear gradient of 4%–28% acetonitrile over 45 min followed by being sprayed into the mass spectrometer with a nanoFlex ion source (Thermo Fisher Scientific). Full-scan MS spectra (300–1,650 *m*/*z*) were acquired in a profile mode with resolution of 60,000 at *m*/*z* 200, a maximum injection time of 20 ms and an automatic gain control (AGC) target value of 3 × 10^6^. Using a 1.4-Th window, up to 15 of the most intense peptides per full scan were isolated for fragmentation by higher energy collisional dissociation with normalized collision energy of 28. MS/MS spectra were acquired at a resolution of 30,000, a maximum injection time of 45 ms and an AGC target value of 1 × 10^5^ in centroid mode. Ions with unassigned charge states, single charged ions, and ions with a charge state of more than four were excluded for fragmentation. To minimize the acquisition of fragment spectra representing already acquired precursors, dynamic exclusion was set to 20 s.

***MS data processing***

MaxQuant (v1.6.17.0) was used to process MS raw data. Acquired spectra were searched against the human ‘one sequence per gene’ database (Taxonomy ID 9606) downloaded from UniProt (12 March 2020; 20,531 sequences), and a collection of 244 common contaminants (‘contaminants.fasta’ provided with MaxQuant v1.6.17.0) using the Andromeda search engine integrated into MaxQuant (v1.6.17.0). Protein identifications were obtained by filtering at false discovery rates below 1% for both peptide spectrum matches and proteins using a target–decoy strategy [1]. Protein quantification and data normalization relied on the MaxLFQ algorithm [2]. The MS proteomics data have been deposited to the ProteomeXchange Consortium through the PRIDE partner repository [3] with the dataset identifiers PXD053709 (Fig. S7). For protein assignment, the UniProt human database v2019 including a list of common contaminants was introduced to correlate with spectra. Tryptic specifications and default settings for mass tolerances in MS and MS/MS spectra were applied for searches. The match-between-run feature was used with a time window of 1 min. For further analysis, Perseus (v2.0.7.0) was applied to obtain the proteins with significant changed between groups after filtering contaminants, reverse entries, and proteins that were only identified by a modified peptide.

***Establishment of mCherry-EGFP-RETREG1 inducible cell lines***

mCherry-EGFP-RETREG1 WT and mCherry-EGFP-RETREG1 mtLIR constructs were cloned into pcDNA5/FRT/TO vector (Thermo Fisher Scientific, V652020) by GATEWAY technology, followed by co-transfection with recombinase pOG44 (Thermo Fisher Scientific, V600520) into Flp-In HeLa TREx cells. Then, 400 μg/ml hygromycin and 15 μg/ml blasticidin were used for selection and maintenance of stable cells. Induction was performed with 2 μg/ml doxycycline for 24 h.

***Analysis of OMV production by SEM***

For ultrastructural analyses, liquid cultures were incubated overnight. Circles of membrane filters (pore size 0.1 μm; Whatman Nucleopore, Cytiva 10419504) were punched out of using a biopsy punch. The filters were placed shiny side up on an agar plate, and 2 μl of bacterial cultures were transferred to the center of the filters. The agar plates were incubated overnight at 37°C. The filters were fixed for 15 min in a 1.5-ml reaction tube containing 2.5% glutaraldehyde in PBS. The fixative was removed and washed three times with PBS. An increasing EtOH series was used to dehydrate the samples. This involved 10%, 30%, 50%, 70%, 90% EtOH for 10 min each, with the last three dilution levels applied twice. Subsequently, the samples were incubated in 100% EtOH for 15 min and critical point drying was performed. After the samples were glued onto SEM stubs (custom made, or Electron Microscopy Sciences, 50-192-8185), they were coated with a 10-nm thick gold layer in order to inhibit charging during microscopy. Afterwards, the samples were imaged using a JEOL IT-200 for overview images, and with a Zeiss Auriga 60 for high-resolution images.

***Analysis of OMV production using negative stain TEM***

For high-resolution ultrastructural analyses, *Salmonella* cultures were grown on agar plates, then bacterial material was picked up with an inoculation loop and resuspended in 500 μl PBS. Here, the samples were carefully inverted to inhibit detachment of OMVs. Then, 5 μl of the bacterial suspension was transferred to a glow-discharged EM copper slot grid (with Formvar charcoal coating [Electron Microscopy Sciences, FCF200-Cu]) and the cells were allowed to adhere for 5 min. The suspension was carefully removed with a filter paper and subsequently washed three times with 5 μl MilliQ. For negative stain, 1% uranyl acetate was applied to the grid for 1 min, and the excess UA was then removed with a filter paper. Afterwards, the samples were imaged by the Zeiss LEO912 and the JEOL JEM 2100Plus.

**REFERENCES**

[1]	Elias JE, Gygi SP. Target-decoy search strategy for increased confidence in large-scale protein identifications by mass spectrometry. Nat Methods. 2007;4(3):207-14.

[2]	Cox J, Hein MY, Luber CA, et al. Accurate proteome-wide label-free quantification by delayed normalization and maximal peptide ratio extraction, termed MaxLFQ. Mol Cell Proteomics. 2014;13(9):2513-26.

[3]	Perez-Riverol Y, Bai J, Bandla C, et al. The PRIDE database resources in 2022: a hub for mass spectrometry-based proteomics evidences. Nucleic Acids Res. 2022;50(D1):D543-D552.

**Table S1.** The oligonucleotides for construction of *Salmonella* mutants.

Oligonucleotides (5’ – 3 ‘)

tolA Del13 For:

GAGTAACAGGCGAACAGTTTTTTGGGAACCGAGAGTGTCAATTCCGGGGATCCGTCGACC

tolB Del13 Rev:

TTCTTTAATTCCTTTAGTAATCAATTAATTATTATCACAGTGTAGGCTGGAGCTGCTTCG

tolA DelCheck For:

CGTTAGCGTCCATACCAGTG

pagL Del For:

AAATAACTATTGACATTGAAATGGTGGTGGAGTGTATATGgtgtaggctggagctgcttc

pagL Del Rev:

CTTCAGCCAGCAACTCGCTAATTGTTATTCAACTTCAGAACATATGAATATCCTCCTTAg

pagL DelCheck For:

GTGACATAACAGAAGTGTGG

degS Del13 For:

TGCTGCTGCCGTTCCCTTTTTTAACGACGCCTCCATCATGATTCCGGGGATCCGTCGACC

degS Del13 Rev:

CTAGTCGTCGAACGACGCGGGCAAAAGCCGCGTCGTTTTATGTAGGCTGGAGCTGCTTCG

degS DelCheck For:

GAAATGCGCAAAGTGATGGC

ompA Del13 For:

ATGGCGTATTTTGGATGATAACGAGGCGCAAAAAATGAAAATTCCGGGGATCCGTCGACC

ompA Del13 Rev:

CCCCGCGACGCGGGGTTTTTTATCAGACGGAAACTTAAGCTGTAGGCTGGAGCTGCTTCG

ompA DelCheck For:

AGCCGTAGATATCGGTAGAG

K1 Red Del:

CAGTCATAGCCGAATAGCCT

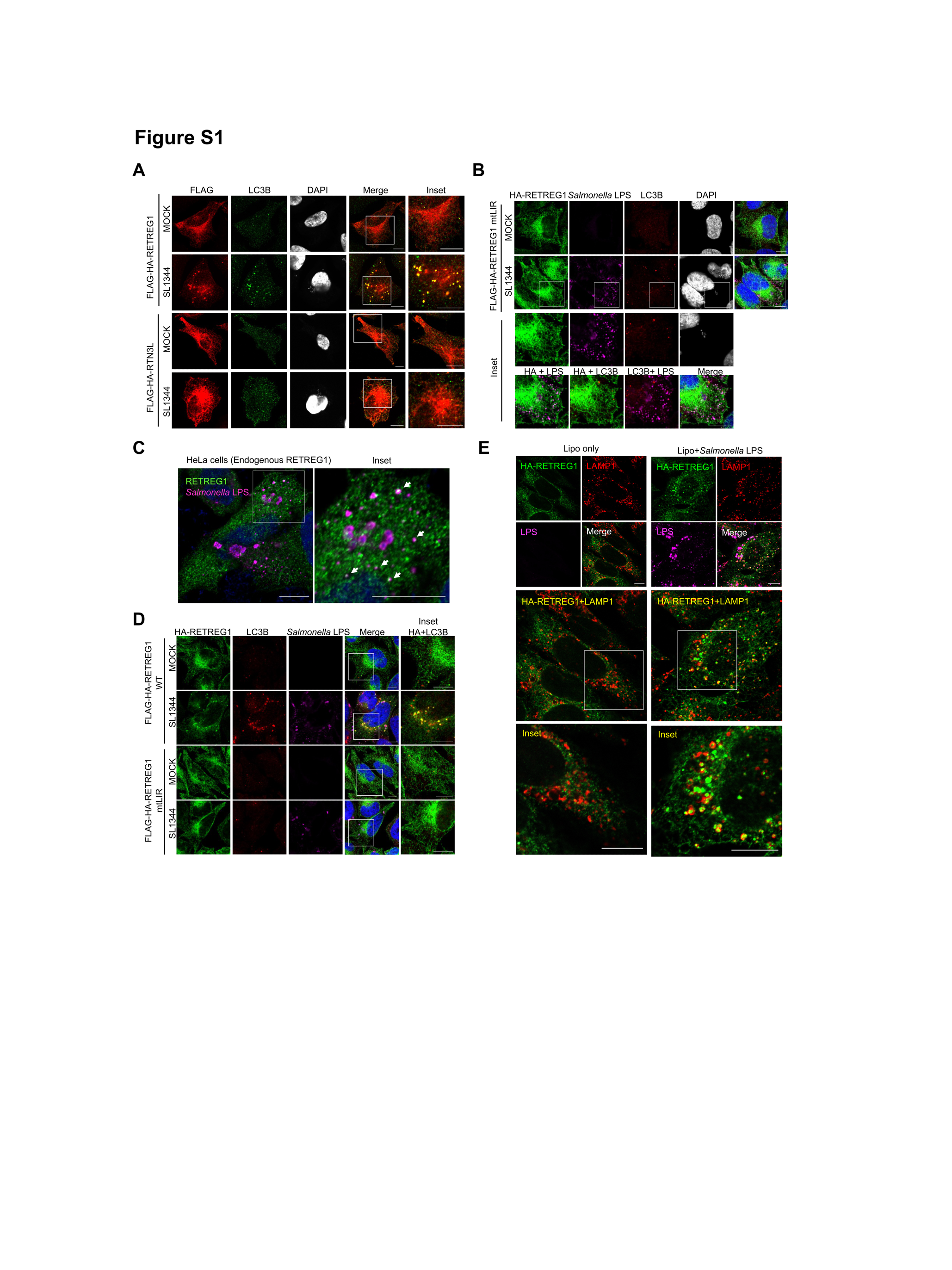

**Figure S1.** *Salmonella* infection mainly activates RETREG1-mediated ER fragmentation. (**A**) Immunofluorescence staining of FLAG and LC3B in HeLa TREx inducible cell lines expressing FLAG-HA-RETREG1 or FLAG-HA-RTN3L, which were infected with SL1344 for 2 h. DAPI was used for cell nuclear and bacterial DNA staining. Scale bar: 10 μm. (**B**) Immunofluorescence staining of HA, *Salmonella* LPS, and LC3B in HeLa TREx inducible cell lines expressing LIR mutant (mtLIR) FLAG-HA-RETREG1 infected with SL1344 for 2 h. DAPI was used for cell nuclear and bacterial DNA staining. Scale bar: 10 μm. (**C**) Immunofluorescence staining of RETREG1 and *Salmonella* LPS in HeLa cells infected by SL1344 for 2 h. Arrows indicate the puncta with colocalization of RETREG1 and *Salmonella* LPS. (**D**) Immunofluorescence staining of HA and LC3B in HeLa TREx inducible cell lines expressing wild type (WT) or mtLIR FLAG-HA-RETREG1 infected with SL1344 for 2 h. DAPI was used for cell nuclear and bacterial DNA staining. Scale bar: 10 μm. (**E**) Immunofluorescence staining of HA, LAMP1, and *Salmonella* LPS in HeLa TREx inducible cell lines expressing FLAG-HA-RETREG1, which were transfected with 1 μg/ml *Salmonella* LPS for 18 h using Lipofectamine 2000. Lipo, lipofectamine only.

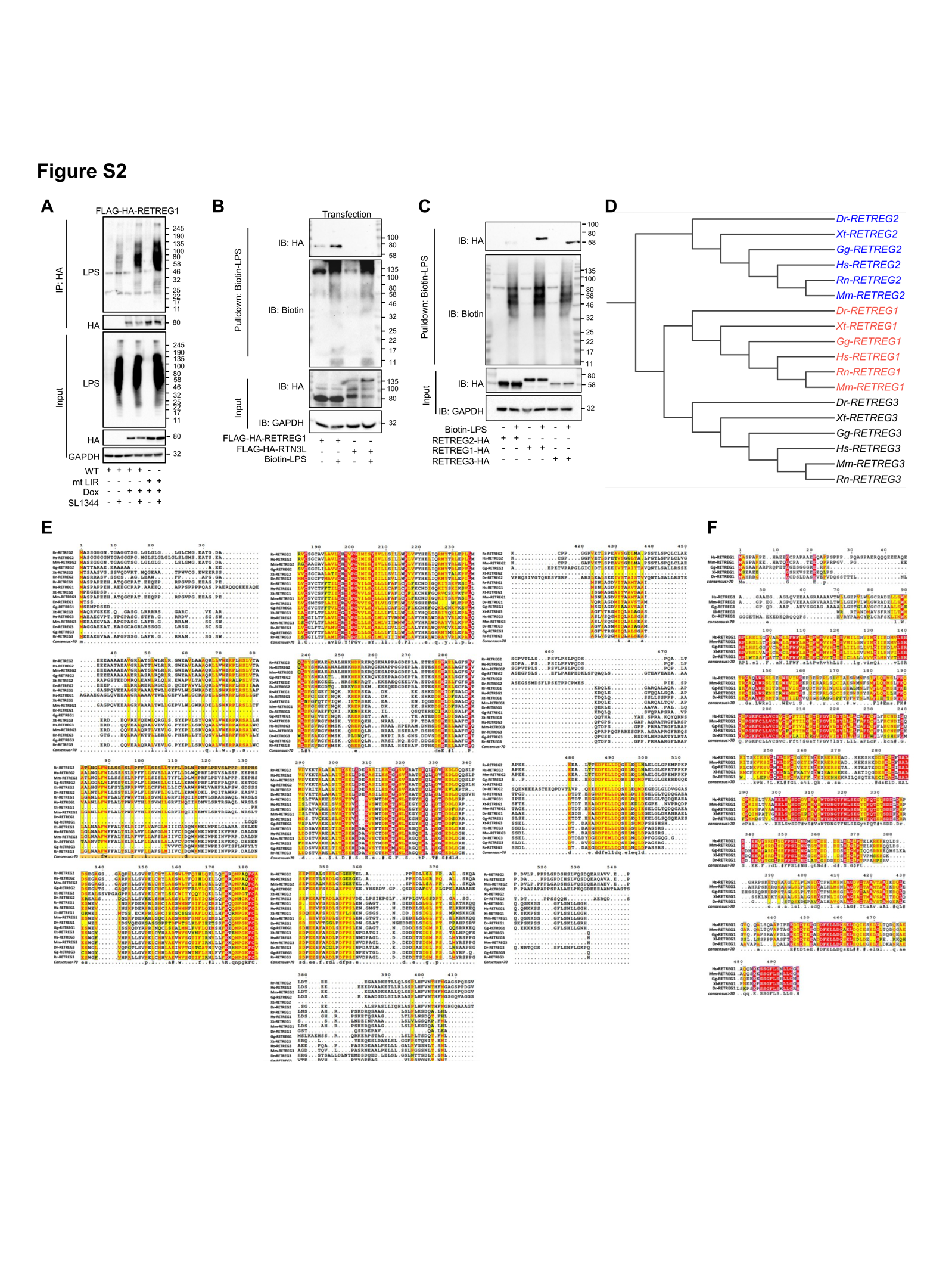

**Figure S2.** LPS bind to RETREG1 and RETREG3, but not RTN3L and RETREG2. (**A**) Co-IP of LPS with HA in HeLa TREx inducible cell lines without Dox treatment or expressing FLAG-HA-RETREG1 WT or FLAG-HA-RETREG1 mtLIR with Dox treatment infected with SL1344 100 moi for 2 h. GAPDH was used as an internal control of input. (**B**) Streptavidin pulldown assays for determining the binding of the transfected biotin-conjugated *E. coli* O111:B4 LPS (10 μg) to HA-tagged RETREG1 or RTN3L in FLAG-HA-RETREG1- or FLAG-HA-RTN3L-expressing HeLa TREx cell lines. GAPDH was used as an internal control of input. (**C**) Streptavidin pulldown assays determining the binding of the biotin-conjugated *E. coli* O111:B4 LPS to HA-tagged RETREG2, RETREG1, or RETREG3 in HA-RETREG2-, HA-RETREG1-, or HA-RETREG3-transfected HeLa cell lysates. GAPDH was used as an internal control of input. (**D**-**F**) Phylogenetic tree of RETREG proteins: Phylogenetic tree of RETREG2, RETREG1 and RETREG3 was generated by using ClustalW followed by analyzing the data through the iTOL server (<https://itol.embl.de/>). Representative sequences were derived from the FASTA for the following organism across phylum Chordata. Dr: *Danio rerio*, Xt: *Xenopus tropicalis*, Gg: *Gallu gallus*, Rn: *Rattus norvegicus*, Mm: *Mus musculus* and Hs: *Homo sapiens*.

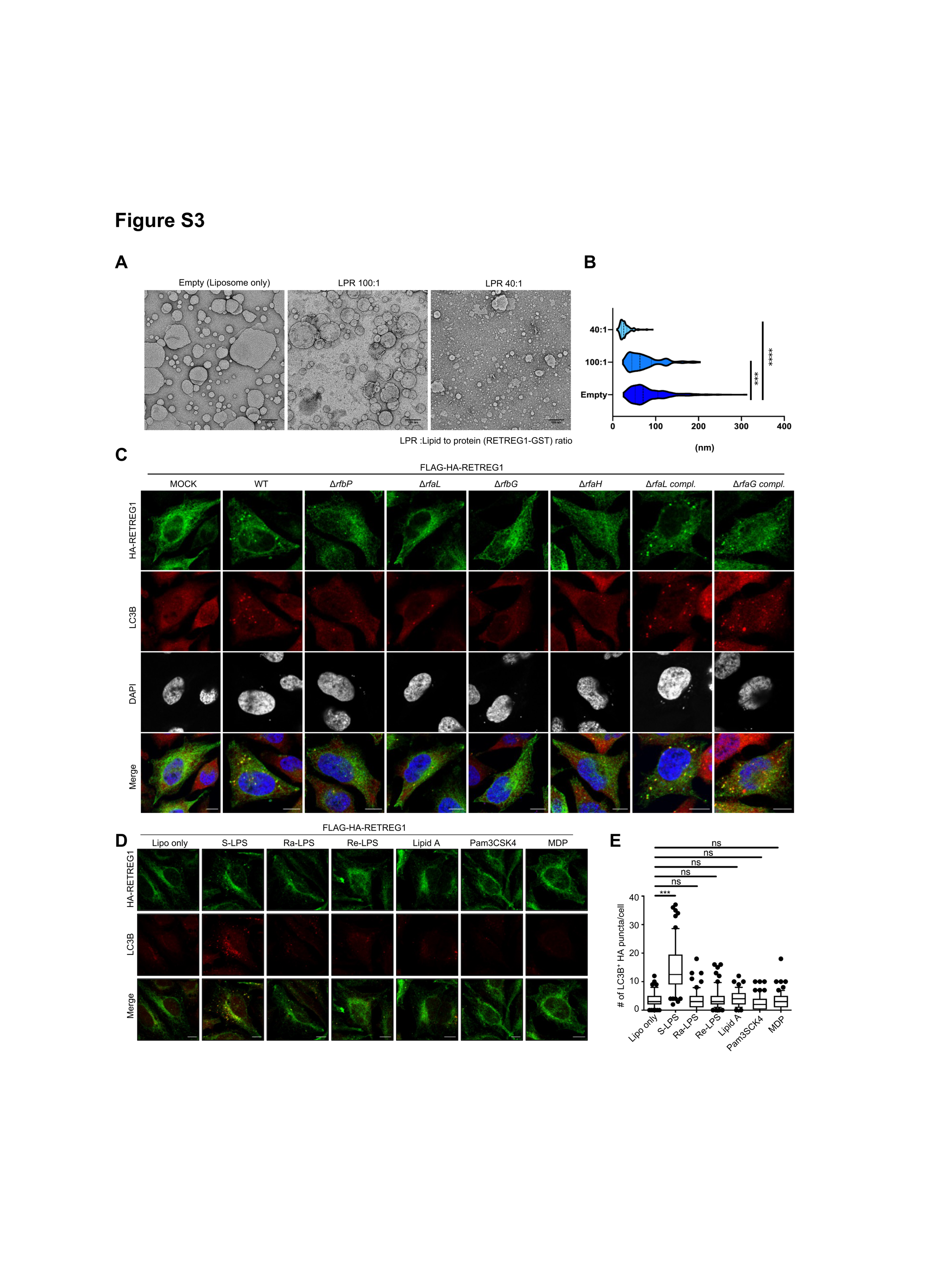

**Figure S3.** S-LPS trigger membrane fragmentation. (**A** and **B**) Representative images (**A**) of remodeled proteoliposomes (scale bars: 100 nm) taken from nsTEM. Empty liposomes (Empty) were added to RETREG1-GST recombinant protein at a lipid to protein ratio (LPR) 100:1 (LPR 100:1) or 40:1 (LPR 40:1). The size distributions (n=300 each) from nsTEM images were measured and presented by violin plots, showing a central boxplot (black lines indicate median with interquartile range) with mirrored histograms on both sides (**B**). Differences were statistically analyzed by one-way ANOVA and Tukey’s multiple comparison test. ***p < 0.001. ns, not significant. (**C**) Immunofluorescence of HA, *Salmonella* LPS, and LC3B in HeLa TREx inducible cell line expressing FLAG-HA-RETREG1 were infected with NCTC12023 WT or LPS O-Ag mutant (Δ) and complemented (compl.) strains for 2 h. DAPI staining was used for cell nuclear and bacterial DNA staining. Scale bar: 10 μm. (**D** and **E**) Immunofluorescence staining of HA and LC3B in HeLa TREx inducible cell line expressing FLAG-HA-RETREG1, transfected with 1 μg/ml S-LPS, Ra-LPS, Re-LPS, Lipid A, Pam3CSK4, and MDP for 18 h by lipofectamine 2000 (**D**). Lipo, lipofectamine only. DAPI was used for cell nuclear DNA staining. Scale bar: 10 μm. Quantification of LC3B-positive HA-RETREG1 puncta per cell (**E**). Solid bars of boxes indicate the medians. Boxes represent interquartile range from 25^th^ to 75^th^ percentile, and whiskers indicate 10^th^ to 90^th^ percentile. Differences were statistically analyzed by one-way ANOVA and Tukey’s multiple comparison test. ***p < 0.001. ns, not significant. Data were collected 60 cells from three independent biological replicates.

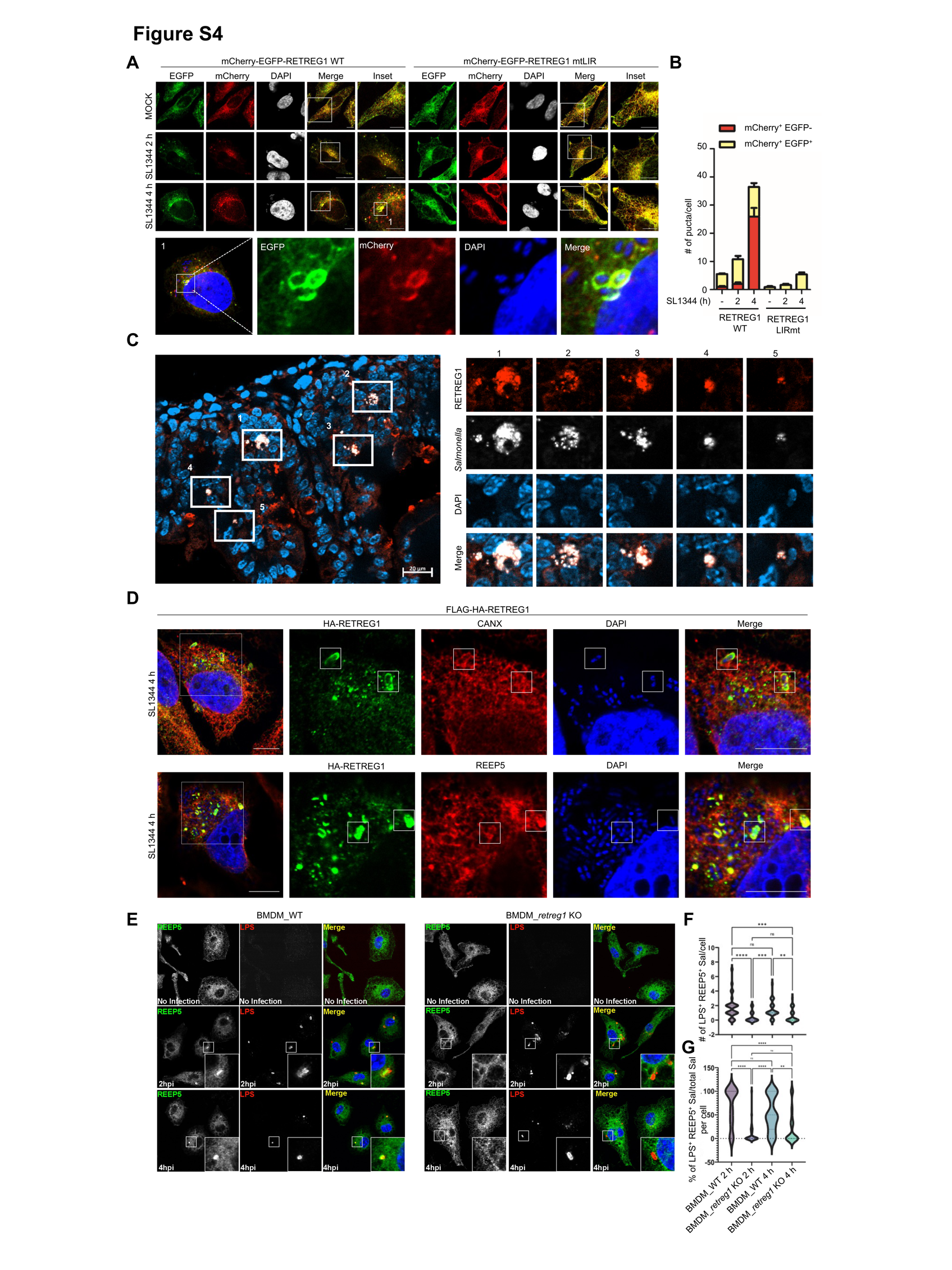

**Figure S4.** *Salmonella* activates RETREG1-mediated ER-phagy and ER fragmentation, forming bacteria-containing vacuoles positive for RETREG1 and ER markers. (**A** and **B**) HeLa TREx inducible cell lines expressing mCherry-EGFP-RETREG1 WT or mCherry-eGFP-RETREG1 mtLIR were infected with SL1344 for 2 h or 4 h (**A**). The inset of square 1 showed the *Salmonella* surrounded by mCherry and EGFP double positive-RETREG1 (**A**). Quantification of mCherry-positive and EGFP-negative puncta (red) and double-positive (yellow) puncta per cell (**B**). DAPI staining was used for cell nuclear and bacterial DNA staining. Scale bar: 10 μm. (**C**) Immunofluorescence staining of RETREG1 (red) in *Salmonella-*infected intestinal tissue section obtained at day 4 post-infection. *Salmonella* GFP staining was used for detecting GFP plasmid carried by *Salmonella* (white). DAPI was used for nuclear DNA staining. Scale bar: 20 μm. (**D**) Immunofluorescence staining of HA and CANX or REEP5 in HeLa TREx inducible cell lines expressing FLAG-HA-RETREG1, which were infected with SL1344 for 4 h. DAPI was used for nuclear and bacterial DNA staining. Scale bar: 10 μm. (**E**-**G**) Immunofluorescence staining of REEP5 (ER marker) and *Salmonella* LPS in WT BMDMs or *retreg*1 KO BMDMs, which were infected with SL1344 for 2 h (2 hpi) or 4 h (4 hpi). *Salmonella* LPS staining was used to indicate bacteria. Quantification of immunofluorescence analysis represented as (**F**) number of vacuoles stained with LPS and REEP5 per cell, (**G**) percentage of *Salmonella* (LPS) positive to REEP5 from total *Salmonella*. Data are mean ± SEM of the number of ER-positive *Salmonella*-containing vacuoles (LPS^+^ and REEP^+^) and % of these vacuoles (LPS^+^ and REEP5^+^) from total *Salmonella* (LPS^+^) per cell (2 h: *n* = 34 for the *retreg*1 KO, *n* = 30 for the WT and 4 h: *n* = 36 for *retreg*1 KO and *n* = 29, Kruskal–Wallis/Dunn’s post-test).  **p<0.01, **p<0.0001, ****p<0.0001 ns, not significant.

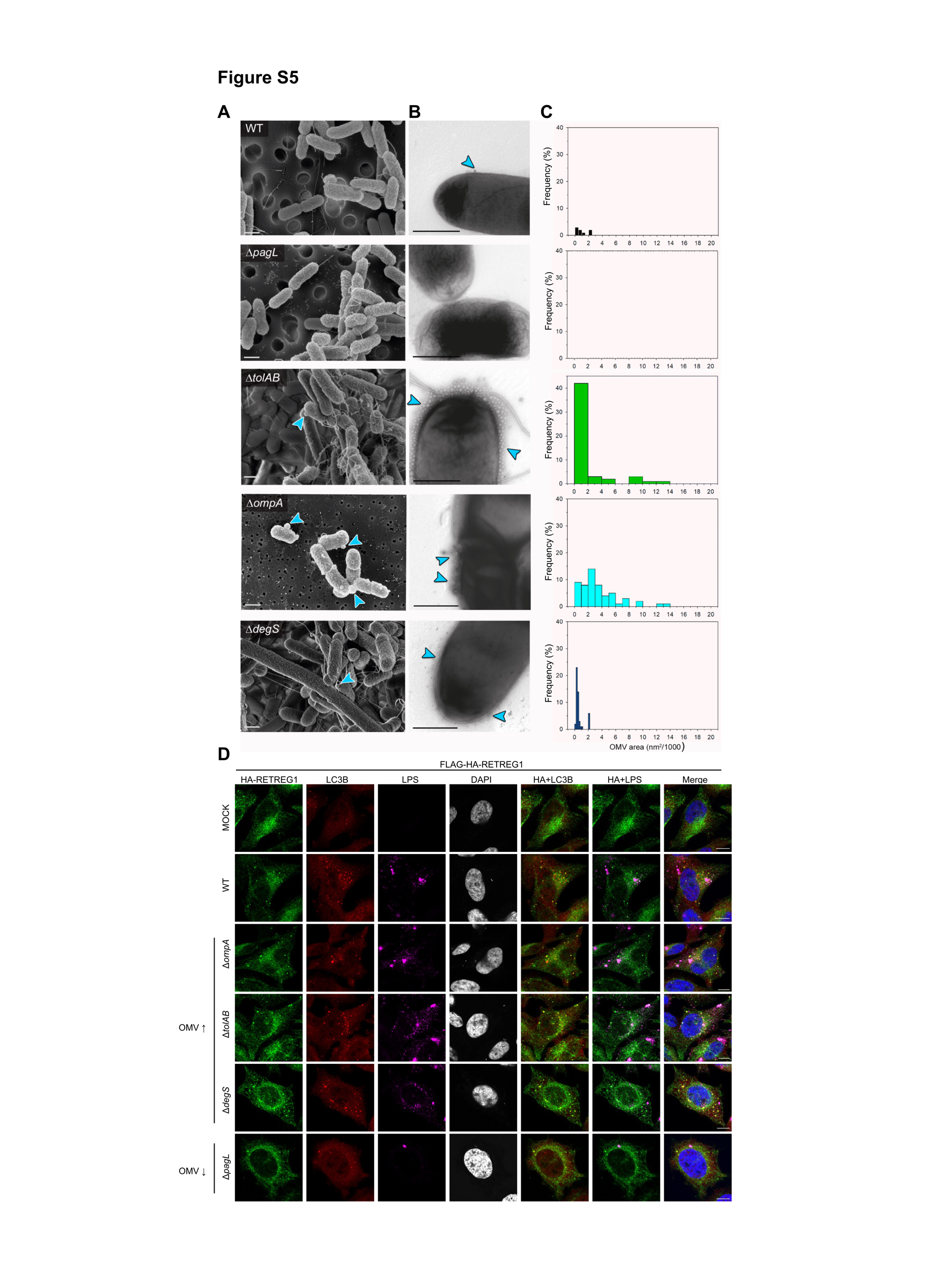

**Figure S5.** Formation of OMV by various *Salmonella* strains and OMV size distribution. (**A**-**C**) STM WT, or mutant strains deficient in *pagL*, *tolAB*, *ompA*, or *degS* were grown on filter paper placed on LB agar (**A**), or overnight on LB agar to allow formation of colonies (**B**). (**A**) Cells were fixed, dehydrated, and coated with 10 nm gold and prepared for SEM imaging on JEOL IT-200 and Zeiss Auriga systems. (**B**) For TEM micrographs, samples were prepared using negative stain. The study was performed as triplicates, 20 bacteria per strain were recorded. Cultures were grown statically on filter papers on LB agar. The experiment was performed as duplicates, at least ten images were generated from each strain. Representative OMV are indicated by arrowheads. Scale bars: 500 nm. (**C**) Size distribution of OMV was determined using morphometric analyses in FIJI. For this, the areas of OMV of at least six bacteria per strains were quantified and the frequency of various OMV size populations was plotted as histogram. The WT strain possessed OMVs of 918 nm^2^ on average, and OMVs up to 2,000 nm^2^ were also detected. No vesicles could be detected for Δ*pagL*. The other three mutant strains showed increased production of OMVs compared to WT strain, and the average sizes of OMVs were 1,611 nm^2^ in Δ*tolAB*, 3,153 nm^2^ in Δ*ompA*, and 590 nm^2^ in Δ*degS*. (**D**) Immunofluorescence staining of HA, *Salmonella* LPS, and LC3B in cells infected with different mutant strains, including Δ*ompA*, Δ*tolAB*, and Δ*degS*, and Δ*pagL* strains. DAPI was used for cell nuclear and bacterial DNA staining. Scale bar: 10 μm.

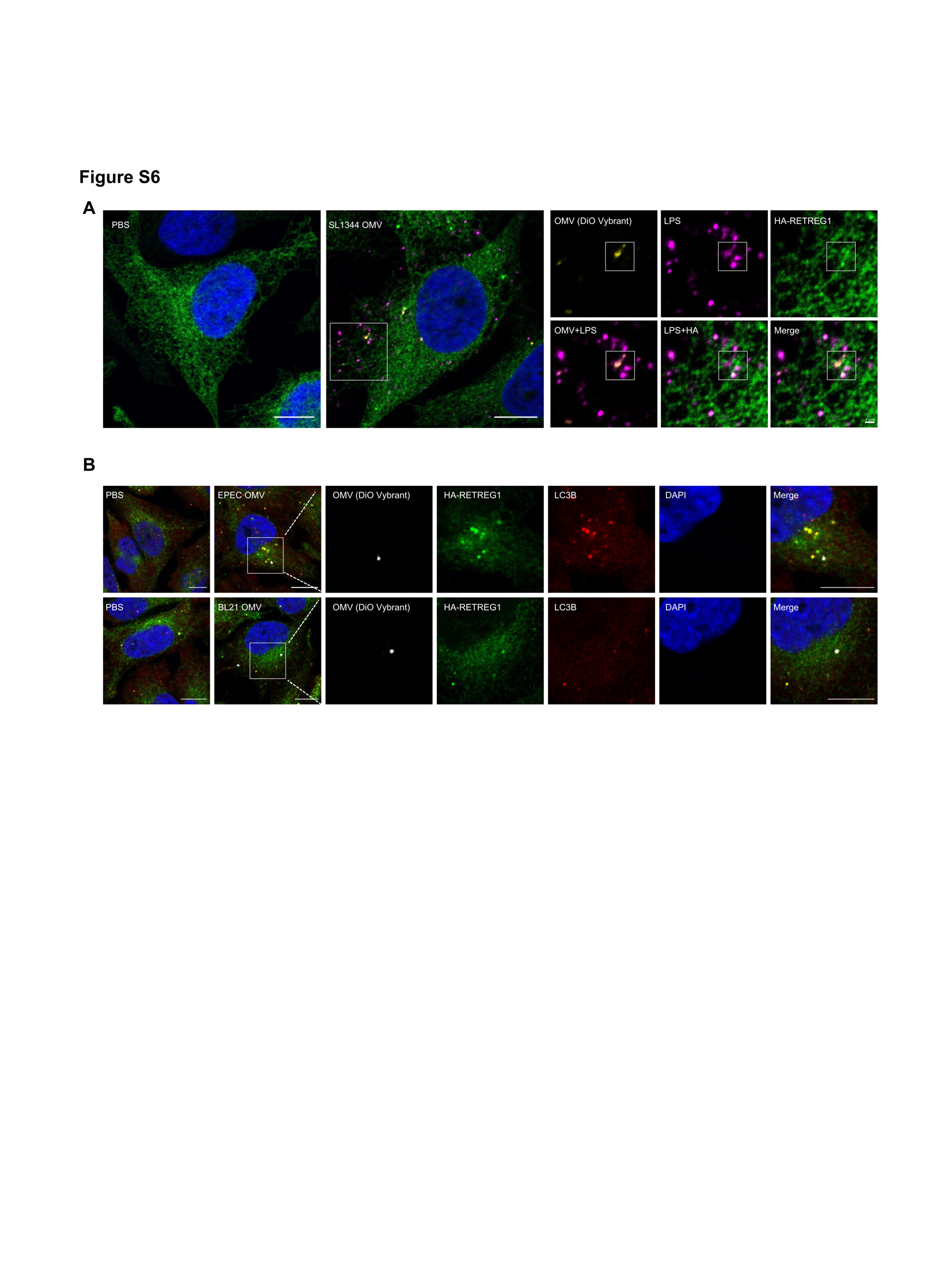

**Figure S6. RETREG1 colocalized with LPS, which is not associated with OMVs.** (**A**) *Salmonella* OMVs were stained by DiO Vybrant dye first, then applied to HeLa TREx inducible cell lines expressing FLAG-HA-RETREG1 for 4 h followed by Immunofluorescence staining of HA and *Salmonella* LPS. DAPI was used for nuclear DNA staining. Scale bar: 10 μm. (**B**) EPEC or BL21 OMVs were stained by DiO Vybrant dye first, then applied to HeLa TREx inducible cell lines expressing FLAG-HA-RETREG1 for 4 h followed by Immunofluorescence staining of HA and LC3B.

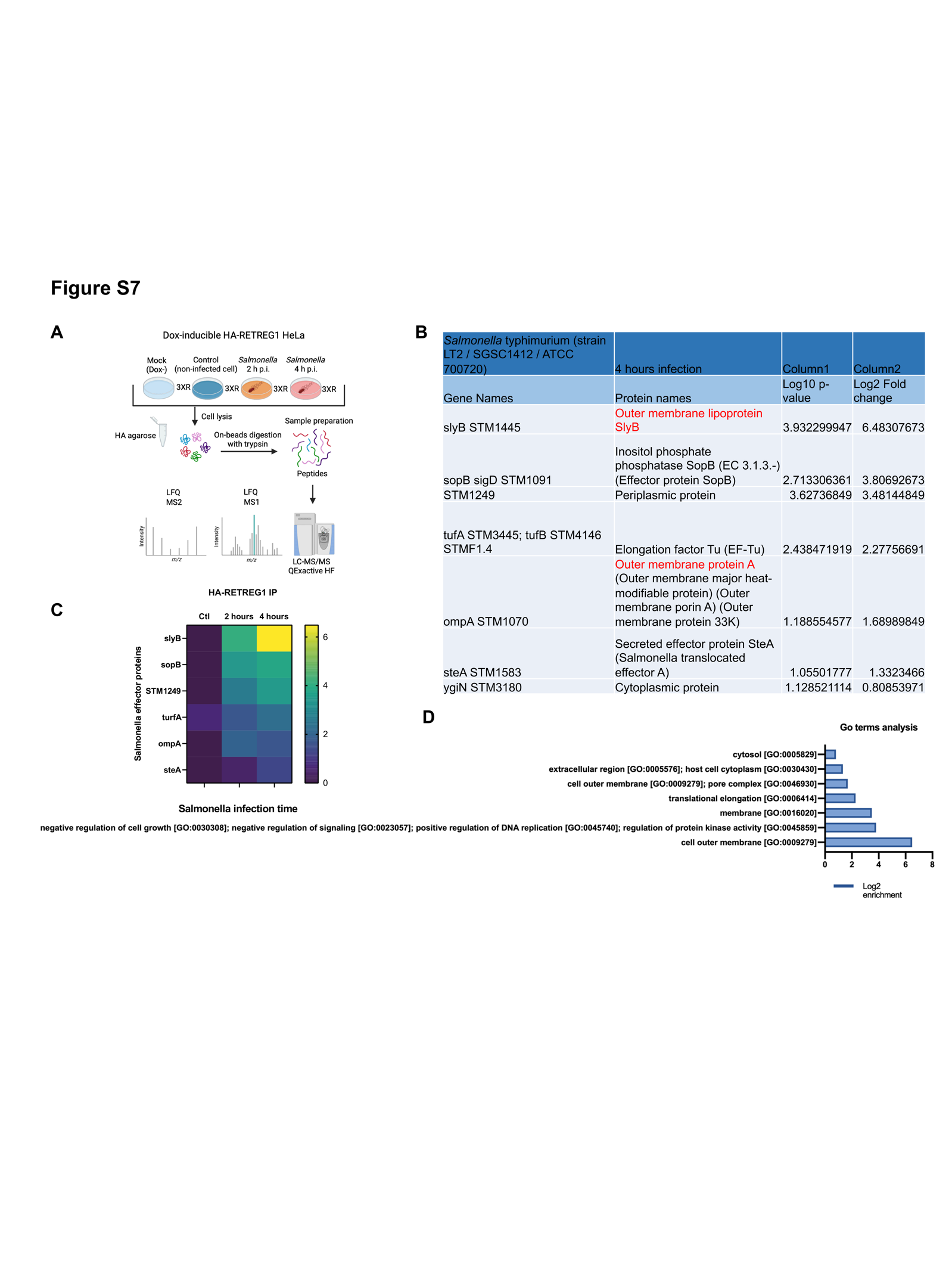

**Figure S7.** RETREG1-interacting *Salmonella* proteins obtained from the interactome of RETREG1. (**A**) MS workflow for the analysis of HA–RETREG1 co-immunoprecipitated from cell lysates following *Salmonella* infection (2 h and 4 h) and Control (non-infected cells, blue) or from mock (HA empty, light blue). HF, Q Exactive HF (Ultra-High-Field Orbitrap) mass spectrometer; LFQ, label free quantification; 3xR 3 biological replicates. (**B**) Table showing the list of *Salmonella* effectors found in the HA-RETREG1 co-immunoprecipitation. Significant and enriched hits are colored in red (significant values were log_2_ enrichment > 1.0 and –log_10_ p value > 1.3). Data are means ± s.d. of *n* = 3 independent experiments. Heat map comparing the interaction of Salmonella effector proteins with WT RETREG1 after 2 or 4 h of infections (immuno-isolated using HA beads). Interaction partners with log_2_ enrichment > 1.0 and –log_10_ p value > 1.3 were plotted. *n* = 3 independent experiments, one-tailed unpaired Student’s test. (**C**) Annotation enrichment analysis of the *Salmonella* effector proteins interacting with RETREG1. Bars represent significantly enriched gene ontology biological process (GOBP), gene ontology cellular component (GOCC) for the identified *Salmonella* proteins.
